# Mutual Regulation between Phosphofructokinase 1 platelet isoform and VEGF Promotes Glioblastoma Tumor Growth

**DOI:** 10.1101/2021.10.14.464342

**Authors:** Je Sun Lim, YuJie Shi, So Mi Jeon, Su Hwan Park, Chuanbao Zhang, Yun-Yong Park, Rui Liu, Jing Li, Wan-Seob Cho, Linyong Du, Jong-Ho Lee

## Abstract

Glioblastoma (GBM) is highly vascular malignant brain tumor that overexpresses vascular endothelial growth factor (VEGF) and phosphofructokinase 1 platelet isoform (PFKP), which catalyzes a rate-limiting reaction in glycolysis. However, it remains unknown whether PFKP and VEGF are reciprocally regulated during GBM tumor growth. Here, we show that PFKP promotes EGFR activation-induced VEGF expression in HIF-1α-dependent and -independent manners in GBM cells. Importantly, we demonstrate that EGFR-phosphorylated PFKP Y64 has critical roles in the AKT/SP1-mediated transcriptional expression of *HIF-1α* and in the AKT-mediated β-catenin S552 phosphorylation, to fully enhance *VEGF* transcription and subsequent blood vessel formation and brain tumor growth. The levels of PFKP Y64 phosphorylation in human GBM specimens positively correlate with HIF-1α expression, β-catenin S552 phosphorylation, and VEGF expression. Conversely, VEGF upregulates PFKP expression in a PFKP S386 phosphorylation-dependent manner, leading to increased PFK enzyme activity, aerobic glycolysis, and proliferation in GBM cells. These findings highlight a novel mechanism underlying the mutual regulation that occurs between PFKP and VEGF for promoting GBM tumor growth and provide the therapeutic potential of targeting the PFKP/VEGF regulatory loop for GBM patients.

## Introduction

Angiogenesis is indispensable for both physiologic and pathologic processes (Apte *et al*, 2019; Hanahan & Weinberg, 2011) and vascular endothelial growth factor (VEGF) is a critical regulator of both processes (Apte *et al*., 2019; Hanahan & Weinberg, 2011). In particular, tumor growth and metastasis highly depend on angiogenesis because the formation of new blood vessels is required for the continued growth and malignant dissemination of solid tumors (Apte *et al*., 2019; Hanahan & Weinberg, 2011). Consequently, excessive angiogenesis is often observed in the pathogenesis of most tumors (Hanahan & Weinberg, 2011). Glioblastoma (GBM) is the most frequent primary adult brain tumor with highly-developed abnormal vascular structures and VEGF overexpression (Jain *et al*, 2007). Defining the mechanisms that regulate VEGF expression in GBM cells has important implications for understanding tumor progression, thereby providing clinically relevant information that may suggest strategies for blocking angiogenesis in the pathogenesis of GBM tumor.

Hypoxia is a common feature of the microenvironment of solid tumors (Pouyssegur *et al*, 2006), and is a well-known stimulus for upregulating VEGF *via* the transcription factor hypoxia-inducible factor-1 (HIF-1) (Semenza, 1998; Shweiki *et al*, 1992). HIF-1, which is a heterodimeric protein consisting of an α subunit that is induced by hypoxia and a β subunit that is constitutively present (Semenza, 1998), binds to a specific consensus sequence within the hypoxia response element (HRE) in the VEGF promoter (Forsythe *et al*, 1996). However, many cancer cells, including GBM cells, express high levels of VEGF, even under normoxic conditions (Feldkamp *et al*, 1999; Hung *et al*, 2016; Maity *et al*, 2000; Pore *et al*, 2004; Ravindranath *et al*, 2001), suggesting that intrinsic factors lead to VEGF upregulation independent of the environment. Epidermal growth factor receptor (EGFR) is one of the elements, which is overexpressed or mutated in many types of cancers, including GBM (Cancer Genome Atlas Research, 2008), and correlates with poor clinical prognosis (Avraham & Yarden, 2011). Although EGFR activation has been shown to upregulate VEGF expression in many types of cancers (Hung *et al*., 2016; Maity *et al*., 2000; Ravindranath *et al*., 2001), the specific mechanisms remain to be elucidated.

Metabolic reprogramming is an emerging hallmark of cancer (Hanahan & Weinberg, 2011). Metabolic reprogramming and other cellular activities are essential for biological processes which can be mutually regulated and are important for tumor development (Xu *et al*, 2021). Beside their canonical metabolic functions, metabolic enzymes possess nonmetabolic functions in cancer cells, which play crucial roles in tumorigenesis (Xu *et al*., 2021). In the glycolytic pathway, phosphofructokinase 1 (PFK1) catalyzes the conversion of fructose 6-phosphate and ATP to fructose-1,6-bisphosphate and ADP, which is one of the key regulatory and rate-limiting steps of glycolysis (Mor *et al*, 2011). PFK1 exists in multiple tertrameric isozymic forms consisting of three types of isoforms: platelet isoform (PFKP), liver isoform (PFKL), and muscle isoform (PFKM); with their expression and composition of isoforms varying depending on the tissue and cell type (Mor *et al*., 2011; Moreno-Sanchez *et al*, 2007). We previously reported that all three isoforms are expressed in GBM cells; PFKP is the prominent PFK1 isoform in GBM cells and is overexpressed in human GBM specimens (Lee *et al*, 2017). Interestingly, we also found that PFKP has a nonmetabolic function; PFKP binds to EGFR upon EGFR activation, leading to EGFR-mediated phosphorylation of PFKP Y64, which in turn binds to an SH2 domain of p85 subunit of phosphoinositide 3-kinase (PI3K) to activate PI3K. The PFKP Y64 phosphorylation-dependent activation of PI3K/AKT enhances aerobic glycolysis in brain tumorigenesis (Lee *et al*, 2018). However, whether PFKP has a critical role in VEGF expression-induced angiogenesis and vice versa during GBM development remains unknown.

In this study, we demonstrate that PFKP promotes EGFR activation-induced VEGF expression in HIF-1α-dependent and -independent manners in GBM cells, leading to enhanced blood vessel formation and brain tumor growth. Conversely, VEGF upregulates PFKP expression, thereby enhancing PFK enzyme activity, aerobic glycolysis, and proliferation in GBM cells.

## Results

### PFKP depletion in GBM cells results in impaired EGFR activation-induced VEGF expression *in vitro* and angiogenesis *in vivo*

To determine whether PFKP expression is required for the angiogenesis-mediated continued growth of GBM tumors, we depleted PFKP expression using PFKP short hairpin RNA (shRNA) in U87 human GBM cells overexpressing constitutively active EGFRvIII mutant (U87/EGFRvIII) (Fig EV1A), which lacks 267 amino acids from its extracellular domain and is frequently found in GBM (Cancer Genome Atlas Research, 2008), and intracranially injected the cells into mice. After implantation and growth, we excised the tumors and performed histologic staining. Depletion of PFKP successfully inhibited the growth of brain tumors derived from intracranially injected U87/EGFRvIII cells at day 18 (Fig 1A). Interestingly, a reduction in PFKP expression resulted in decreased blood vessel formation as evidenced by the intensity of CD31 expression (Fig 1B), in which the tumor sizes at an early stage (day 5) were comparable between control shRNA-expressing and PFKP shRNA-expressing U87/EGFRvIII cells (Fig 1A). These results suggest that PFKP plays an important role in EGFR activation-induced GBM angiogenesis.

**Figure 1.**
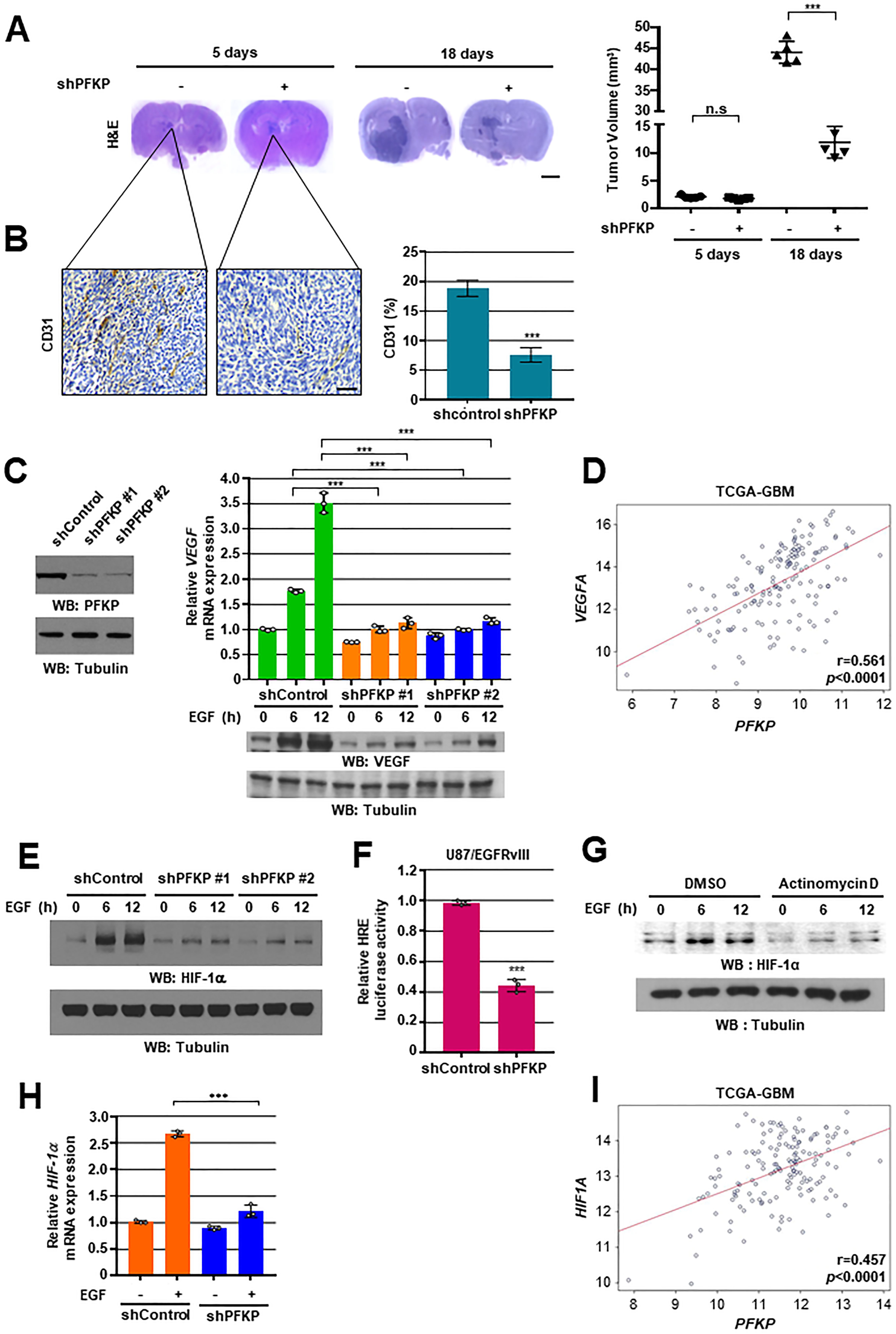
PFKP expression is required for EGFR activation-induced VEGF expression *in vitro* and GBM angiogenesis *in vivo.* WB and qRT-PCR were performed with the indicated primers and antibodies, respectively (**C, E, G, H**). **A** and **B** Representative H&E staining images of intracranial xenografts bearing U87/EGFRvIII cells stably expressing with or without PFKP depletion (**A**; left panel) and quantification of tumor volumes (**A**; right panel). IHC analyses of the tumor tissues with an anti-CD31 antibody (**B**; left panel). Quantification of CD31 (**B**; right panel). Scale bar, 2 mm (**A**) and 100 μm (**B**). **C** U87/EGFR cells with different shRNAs against PFKP (left panel). Serum-starved U87/EGFR cells with or without PFKP depletion were treated with EGF (100 ng/mL) (right panel). **D** TCGA analysis of *PFKP* and *VEGF* mRNA expression from TCGA-GBM data (n = 154). **E** Serum-starved U87/EGFR cells with or without PFKP depletion were treated with EGF (100 ng/mL). **F** HRE luciferase activity was measured in U87/EGFRvIII cells with stable expression of control shRNA or PFKP shRNA. **G** Serum-starved U87/EGFR cells were pretreated with DMSO or actinomycin D (1 μg/mL) for 1 h and then stimulated with or without EGF (100 ng/mL). **H** Serum-starved U87/EGFR cells stably expressing control shRNA or PFKP shRNA were treated with or without EGF (100 ng/mL) for 12 h. **I** TCGA analysis of *PFKP* and *HIF-1α* mRNA expression from TCGA-GBM data (n = 154). The data represent the mean ± SD of three independent experiments (**C, F, H**). \*\*\**P* < 0.001, based on the Student’s t-test.

We next determined the effect of PFKP on EGFR activation-induced VEGF expression *in vitro*. A reduction in PFKP expression largely reduced EGFR activation-induced *VEGF* mRNA expression and its protein expression in U87/EGFR (Fig 1C), U87/EGFRvIII (Fig EV1B), and U251 cells (Fig EV1C). In addition, to determine the clinical relevance of PFKP-regulated VEGF expression, we analyzed The Cancer Genome Atlas (TCGA) data and revealed that the expression levels of *PFKP* were positively correlated with *VEGF* mRNA expression levels in GBMs (Fig 1D) and BRCA-Metabric (breast cancer) (Fig EV1D). Our *in vitro* and *in vivo* results suggest that PFKP plays a critical role in EGFR activation-induced VEGF expression in GBM cells, thereby promoting GBM angiogenesis.

### PFKP expression is required for EGFR activation-induced HIF-1α expression and transactivation

Accumulated evidence has shown that EGFR activation increases the expression of HIF-1α under normoxic conditions in cancer cells (Laughner *et al*, 2001; Maity *et al*., 2000; Phillips *et al*, 2005). To examine the effect of PFKP on EGFR activation-induced HIF-1α expression, we treated EGF to cancer cells, including EGFR-overexpressing U87/EGFR and U251 GBM cells, MDA-MB-231 human breast carcinoma cells, and A431 human epidermoid carcinoma cells, with or without expression of PFKP shRNA. Immunoblotting analysis showed that EGFR activation-induced HIF-1α expression was largely inhibited by depletion of PFKP in U87/EGFR (Fig 1E), U251, MDA-MB-231, A431 cells (Fig EV2A), and U87/EGFRvIII cells (Fig EV2B). In line with these findings, HRE luciferase reporter analysis showed that depletion of PFKP expression largely inhibited EGFR activation-induced HIF-1α transactivation in U87/EGFRvIII cells (Fig 1F). Pretreatment of cancer cells with actinomycin D, a transcription inhibitor, almost abolished EGF-enhanced HIF-1α expression in U87/EGFR (Fig 1G), U251, MDA-MB-231, and A431 cells (Fig EV2C), suggesting that transcriptional regulation of HIF-1α is essentially involved in the response to EGFR activation. Quantitative PCR analyses showed that EGF treatment enhanced mRNA levels of the *HIF-1α* gene and this increase was attenuated by PFKP depletion in U87/EGFR (Fig 1H), U251, MDA-MB-231, and A431 cells (Fig EV2D). Consistent with these findings, depletion of PFKP in U87/EGFRvIII cells resulted in decreased *HIF-1α* mRNA expression (Fig EV2E). Analyses of TCGA data showed that the expression levels of *PFKP* were positively correlated with *HIF-1α* mRNA expression levels in GBMs (Fig 1I) and BRCA-Metabric (breast cancer) (Fig EV2F). Taken together, these results indicate that PFKP expression is required for EGFR activation-induced *HIF-1α* transcriptional expression and its activity in cancer cells.

### PFKP Y64 phosphorylation induces EGFR activation-enhanced *HIF-1α* transcriptional expression through SP1 transactivation

To determine how *HIF-1α* transcriptional expression is regulated by EGFR activation, we pretreated U87/EGFR cells with inhibitors of early signal pathways, including ERK, JNK, p38, PI3K, or NF-κB, which successfully blocked EGF-induced ERK, c-Jun, p38, AKT, or IκBα phosphorylation, respectively (Fig EV3A). Inhibition of PI3K/AKT or NF-κB, but not of ERK, p38, or JNK, largely abrogated EGF-induced HIF-1α protein expression in U87/EGFR cells (Fig 2A). In addition, pretreatment with PI3K inhibitor or NF-κB inhibitor blocked EGF-induced *HIF-1α* mRNA expression in U87/EGFR cells (Fig 2B). In line with this result, pretreatment of U87/EGFR, U251, MDA-MB-231, and A431 cells with MK-2206, a selective AKT1/2/3 inhibitor, blocked EGF-stimulated *HIF-1α* mRNA (Fig 2C and EV3B) and its protein expression (Fig 2D and EV3B). These results indicate that PI3K/AKT and NF-κB pathways are primarily involved in regulation of the EGFR activation-induced *HIF-1α* transcriptional expression in cancer cells. In a previous report, we found that EGFR-phosphorylated PFKP Y64 promotes PI3K/AKT activation (Lee *et al*., 2018). In addition, depletion of PFKP did not alter the other EGFR activation-induced early signals, including phosphorylation of EGFR, ERK, p38, JNK/c-Jun, IκBα, and p65 in U87/EGFR cells (Fig EV3C). Thus, we excluded NF-κB signal for the role of PFKP in EGFR activation-induced *HIF-1α* transcriptional expression. We next investigated the effect of PFKP Y64 phosphorylation on *HIF-1α* mRNA expression in response to EGFR activation. As expected, depletion of endogenous PFKP resulted in decreased *HIF-1α* mRNA (Fig 2E) and protein (Fig 2F) expression in U87/EGFRvIII cells; this decrease was rescued by reconstituted expression of RNAi-resistant (r) WT Flag-rPFKP, but not rPFKP Y64F mutant, (Fig 2E and 2F). Of note, the inhibitory effect of rPFKP Y64F expression on *HIF-1α* mRNA (Fig 2E) and protein expression (Fig 2F) in U87/EGFRvIII cells was restored by expression of the constitutively active AKT1 mutant (HA-myr-AKT1). These results indicate that PFKP Y64 phosphorylation enhances EGFR activation-induced *HIF-1α* mRNA and its protein expression in an AKT activation-dependent manner.

**Figure 2.**
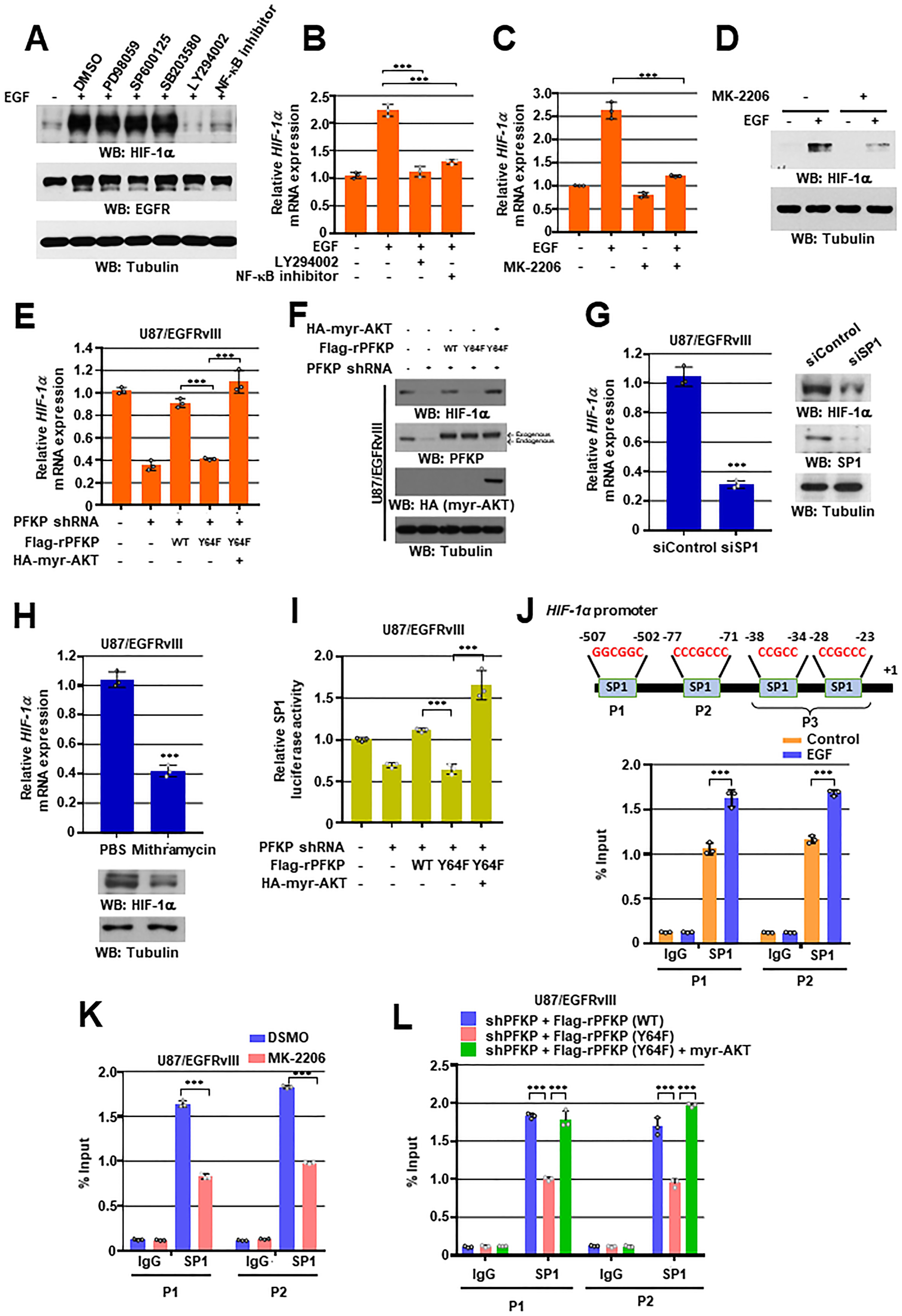
PFKP Y64 phosphorylation induces EGFR activation-enhanced *HIF-1α* transcriptional expression through SP1 transactivation. WB and qRT-PCR were performed with the indicated primers and antibodies, respectively (**A-H**). **A** Serum-starved U87/EGFR cells were pretreated DMSO, PD98059 (20 μM), SP600125 (25 μM), SB203580 (10 μM), LY294002 (20 μM), or NF-κB inhibitor (1 μM) for 1 h and then stimulated with or without EGF (100 ng/mL) for 12 h. **B-D** Serum-starved U87/EGFR cells were pretreated DMSO, LY294002, NF-κB inhibitor (**B**), or MK-2206 (5 μM) (**C, D**) for 1h and then stimulated with or without EGF (100 ng/mL) for 12 h. **E** and **F** U87/EGFRvIII cells with or without PFKP depletion and with or without reconstituted expression of WT Flag-rPFKP or Flag-rPFKP Y64F mutant in the presence or absence of HA-myr-AKT expression. **G** U87/EGFRvIII cells were transfected with control siRNA or SP1 siRNA. **H** U87/EGFRvIII cells were treated with PBS or mithramycin (500 nM) for 12 h. **I** SP1 luciferase activity was measured in U87/EGFRvIII cells with or without PFKP depletion and with or without reconstituted expression of WT Flag-rPFKP or Flag-PFKP Y64F mutant in the presence or absence of HA-myr-AKT expression. **J-L** ChIP assays were performed with anti-SP1 antibody, and real-time PCR analyses were performed with primers against the *HIF-1α* promoter. (**J)** The schematic of the putative SP1 binding site (Marked as P1 – P3) on the *HIF-1α* promoter region (**J**; upper panel). U87/EGFR cells were treated with or without EGF (100 ng/mL) for 12 h (**J**; bottom panel). (**K**) U87/EGFRvIII cells were pretreated DMOS or MK-2206 (5 μM) for 1h and then EGF (100 ng/ml) for 12h. (**L**) U87/EGFRvIII cells with PFKP depletion and with or without reconstituted expression of WT Flag-rPFKP or Flag-rPFKP Y64F mutant were transfected with or without HA-myr-AKT expression. The data represent the mean ± SD of three independent experiments (**B, C, E, G, H-L**). \*\*\**P* < 0.001, based on the Student’s t-test or one-way ANOVA with Tukey’s post hoc test.

It has been reported that the transcription factor specific protein 1 (SP1) has putative binding sites on the *HIF-1α* gene promoter (Iyer *et al*, 1998), but, its role and regulatory mechanism in EGFR activation-induced *HIF-1α* transcription are unknown. Depletion of SP1 in U87/EGFRvIII cells (Fig 2G) or treatment with the SP1 inhibitor mithramycin in U87/EGFRvIII (Fig 2H), U87/EGFR, MDA-MB-231, A431 cells (Fig EV3D and EV3E) resulted in reduced EGFR activation-induced *HIF-1α* mRNA and protein expression levels. These data indicate that SP1 is required for *HIF-1α* transcriptional expression in response to EGFR activation. PI3K/AKT pathway is associated with increased phosphorylation of SP1 and its transcriptional activity (Chuang *et al*, 2013; Pore *et al*., 2004; Reisinger *et al*, 2003; Sroka *et al*, 2007; Tan & Khachigian, 2009). As PFKP Y64 phosphorylation regulates PI3K/AKT in EGFR-activated cancer cells, we investigated whether PFKP Y64 phosphorylation could modulate SP1 activity. Luciferase reporter analysis using a plasmid containing the luciferase reporter gene driven by SP1-responsive promoter showed that the transcriptional activity of SP1 in the U87/EGFRvIII cells expressing rPFKP Y64F in PFKP- depleted U87/EGFRvIII cells was reduced compared with the cells expressing its WT counterpart, and this reduction was abrogated by expression of an active AKT1 mutant (Fig 2I). Next, we performed a chromatin immune precipitation (ChIP) assay with an anti-SP1 antibody to identify the exact binding site of SP1 in the promoter region of the *HIF-1α* gene in response to EGFR activation. Based on the previous report (Iyer *et al*., 1998), we designed three sets of primers that specifically amplify the indicated regions (P1 for -507 bp to -502 bp; P2 for -77 bp to -71 bp; and P3 for -38 bp to -23 bp, +1 indicates the first bp of exon 1) of the *HIF-1α* gene promoter (Fig 2J; top panel). As shown in Fig 2J (bottom panel), only the P1 and P2 regions were amplified, indicating that SP1 specifically binds to the P1 and P2 regions of the *HIF-1α* gene promoter in response to EGFR activation. MK-2206 treatment reduced the binding of SP1 to the promoter region of *HIF-1α* in U87/EGFR cells (Fig 2K). Furthermore, the binding of SP1 to the promoter region of *HIF-1α* in the U87/EGFRvIII cells expressing rPFKP Y64F in PFKP-depleted U87/EGFRvIII cells was reduced compared with the cells expressing its WT counterpart, and this reduction was alleviated by expression of an active AKT1 mutant (Fig 2L). These results strongly suggest that the PFKP Y64 phosphorylation regulates EGFR activation-enhanced *HIF-1α* transcriptional expression through AKT/SP1 pathway in cancer cells.

### PFKP Y64 phosphorylation induces VEGF expression through HIF-1 α expression and β-catenin Ser552 phosphorylation in response to EGFR activation

We next determined the role of PFKP Y64 phosphorylation in EGFR activation-induced VEGF expression. Depletion of PFKP resulted in decreased EGFR activation-induced *VEGF* mRNA and protein expression levels in U87/EGFRvIII cells (Fig 3A). This decrease was rescued by reconstituted expression of WT Flag-rPFKP, but not the rPFKP Y64F mutant (Fig 3A), suggesting a crucial role of PFKP Y64 phosphorylation in the EGFR activation-induced VEGF expression. Interestingly, the inhibitory effect of rPFKP Y64F on VEGF expression in U87/EGFRvIII cells was not fully rescued by exogenous expression of HIF-1α (Fig 3A). In addition, EGFR activation still induced VEGF expression in U87/EGFR (Fig 3B) and U251 cells (Fig EV4B) even when we blocked HIF-1α transactivation by treatment with a HIF-1α inhibitor (Fig EV4A and EV4C). These results suggest that PFKP Y64 phosphorylation upregulates VEGF expression in HIF-1α-dependent and -independent mechanisms in response to EGFR activation.

**Figure 3.**
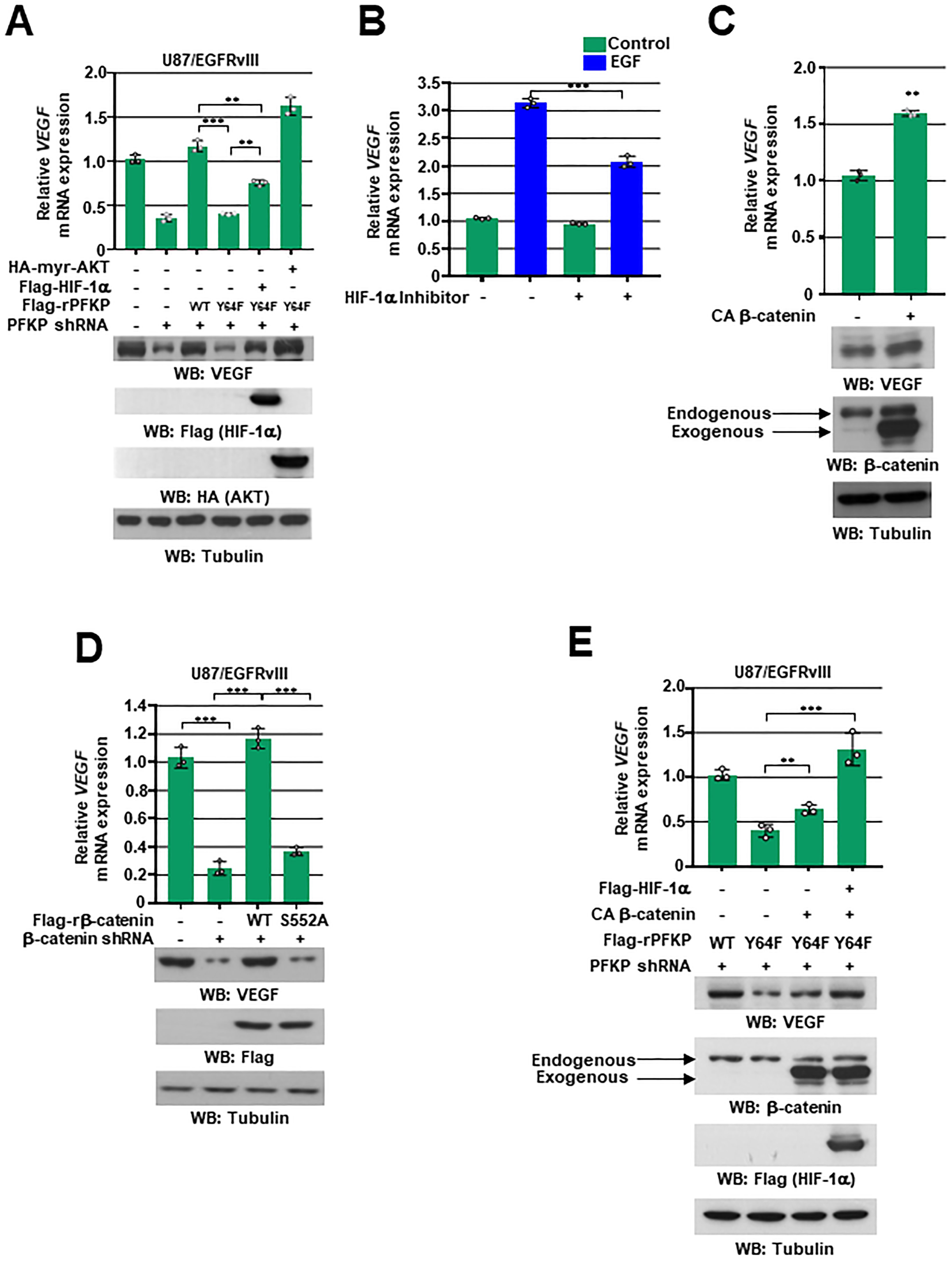
PFKP Y64 phosphorylation induces VEGF expression through HIF-1 α expression and β-catenin Ser552 phosphorylation in response to EGFR activation. WB and qRT-PCR were performed with the indicated primers and antibodies, respectively (**A-E**). **A** U87/EGFRvIII cells with or without PFKP depletion and with or without reconstituted expression of WT Flag-rPFKP or Flag-rPFKP Y64F mutant were transfected with or without Flag-HIF-1α and HA-myr-AKT expression. **B** Serum starved U87/EGFR cells were pretreated DMSO or HIF inhibitor (10 μM) for 1h and then treated EGF (100 ng/mL) for 12h. **C** U87/EGFR cells were transfected with control vector or CA β-catenin. **D** U87/EGFRvIII cells with stable expression of the β-catenin shRNA or a control shRNA were reconstituted with or without WT rβ-catenin or rβ-catenin S552A mutant. **E** PFKP-depleted U87/EGFRvIII cells with or without reconstituted expression of WT Flag-rPFKP or Flag-rPFKP Y64F mutant were transfected with or without CA β-catenin and Flag-HIF-1α expression. The data represent the mean ± SD of three independent experiments (**A-E**). \*\**P* < 0.01; \*\*\**P* < 0.001, based on the Student’s t-test or one-way ANOVA with Tukey’s post hoc test.

It has been demonstrated that β-catenin regulates VEGF expression in colon cancer (Easwaran *et al*, 2003). Consistent with this report, expression of the constitutively active β-catenin (CA β-catenin) mutant increased *VEGF* mRNA expression and its protein expression in U87/EGFR cells (Fig 3C). In contrast, depletion of β-catenin reduced *VEGF* mRNA and protein expression levels in U87/EGFRvIII cells (Fig 3D). AKT has been shown to directly phosphorylate β-catenin at S552 to promote the nuclear translocation and activation of β-catenin (Fang *et al*, 2007). Reconstituted expression of WT rβ-catenin, but not the rβ-catenin S552A mutant, successfully restored the reduced VEGF expression levels in the β-catenin-depleted U87/EGFRvIII cells (Fig 3D), suggesting that AKT-dependent β-catenin activation is instrumental for EGFR activation-increased VEGF expression. We previously reported that PFKP induced AKT-mediated β-catenin S552 phosphorylation and subsequent β-catenin transactivation in a PFKP Y64 phosphorylation-dependent manner (Lee *et al*, 2020). Because PFKP Y64F-reduced β-catenin S552 phosphorylation and subsequent β-catenin transactivation is involved in the reduced VEGF expression in PFKP-depleted U87/EGFRvIII cells (Fig 3E; second lane), we ectopically introduced expression of CA β-catenin, and showed that expression of CA β-catenin partially rescued the PFKP Y64F-reduced VEGF expression (Fig 3E; third lane), which was fully rescued by additional exogenous expression of HIF-1α (Fig 3E; fourth lane). In line with this result, expression of an active AKT1 mutant fully rescued the inhibitory effect of PFKP Y64F on VEGF expression (Fig 3A; sixth lane). These results demonstrate that PFKP Y64 phosphorylation plays critical roles in EGFR activation-induced VEGF expression through AKT/SP1-mediated HIF-1α expression and AKT-mediated β-catenin S552 phosphorylation.

### PFKP Y64 phosphorylation induces HIF-1 α expression, β-catenin S552 phosphorylation, and VEGF expression, and promotes blood vessel formation *in vivo*

We next intracranially injected PFKP-depleted U87/EGFRvIII cells with reconstituted expression of WT rPFKP or rPFKP Y64F mutant, with or without an active AKT1 mutant, into athymic nude mice. The growth of brain tumors (Fig 4A), HIF-1α expression, β-catenin S552 phosphorylation, VEGF expression, and blood vessel formation (Fig 4B) in mice which implanted PFKP-depleted U87/EGFRvIII cells expressing rPFKP Y64F were decreased compared with the cells expressing its WT counterpart, and this reduction was restored by expression of an active AKT1 mutant (Fig 4A and 4B). These results indicate that PFKP Y64 phosphorylation-induced AKT activation plays critical roles in EGFR activation-induced HIF-1α expression, β-catenin S552 phosphorylation, VEGF expression, blood vessel formation, and brain tumor growth.

**Figure 4.**
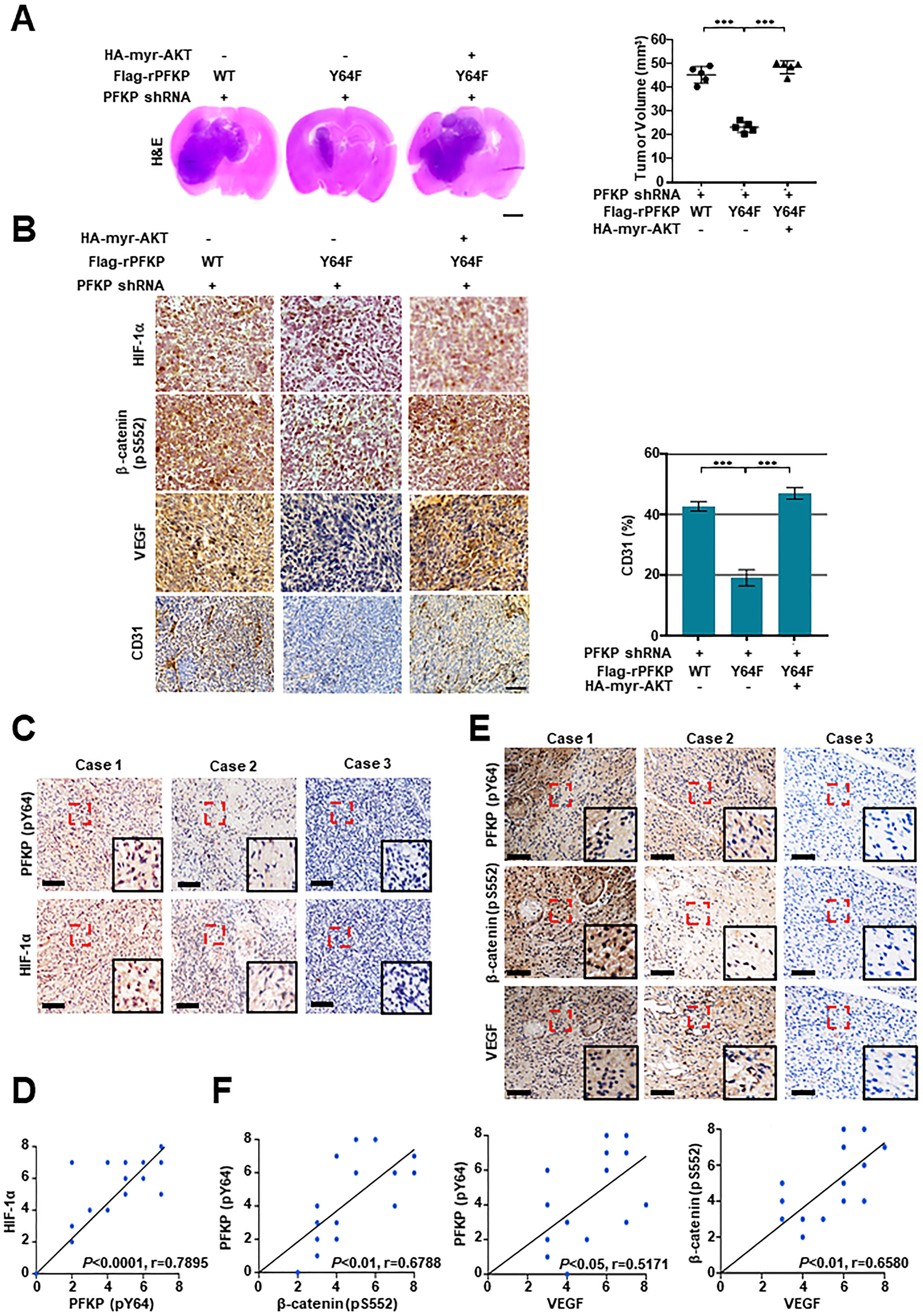
PFKP Y64 phosphorylation induces HIF-1 α expression, β-catenin S552 phosphorylation, and VEGF expression, and promotes blood vessel formation *in vivo*. **A** Representative H&E staining images of intracranial xenografts bearing PFKP-depleted U87/EGFRvIII cells with or without reconstituted expression of WT Flag-rPFKP or Flag-rPFKP Y64F mutant and with or without myr-AKT1 (**A**; left panel) and quantification of tumor volumes (**A**; right panel). Scale bar, 2 mm. **B** IHC analyses of the tumor tissues with the indicated antibodies (**B**; left panel). Quantification of CD31 (**B**; right panel). Scale bar, 100 μm. **C** and **E** IHC staining of human GBM specimens was performed with the indicated antibodies (n = 25). Representative images from the staining of three different specimens are shown. Scale bar, 100 μm. **D** and **F** The IHC stains were scored, and correlation analyses were performed. Pearson correlation test was used. Note that the scores of some samples overlap. \*\*\**P* < 0.001, based on one-way ANOVA with Tukey’s post hoc test.

To determine the clinical significance of PFKP Y64 phosphorylation-mediated HIF-1α expression, β-catenin S552 phosphorylation, and VEGF expression, we analyzed human primary GBM specimens. PFKP Y64 phosphorylation levels were positively correlated with HIF-1α expression, β-catenin S552 phosphorylation, and VEGF expression (Fig 4C and 4E) and these correlations were significant (Fig 4D and 4F).

### VEGF induces PFKP expression, PFK enzyme activity, aerobic glycolysis, and proliferation in GBM cells

We next examined the role and mechanism of angiogenesis-independent functions of VEGF in GBM cells. Interestingly, VEGF stimulation successfully induced glucose consumption (Fig 5A and EV5A), lactate secretion (Fig 5B and EV5B), and proliferation (Fig 5C and EV5C) in U87/EGFR and A172 cells. To investigate which enzymes are regulated during VEGF-enhanced aerobic glycolysis in GBM cells, we analyzed protein expression profiles of glycolytic enzymes using immunoblotting. The protein expression of glycolytic enzymes, including hexokinase 2 (HK2), PFKM, PFK2, Aldolase, glyceraldehyde 3-phosphate dehydrogenase (GAPDH), phosphoglycerate kinase 1 (PGK1), phosphoglycerate mutase 1 (PGAM1), Enolase, pyruvate kinase M2 (PKM2), and lactate dehydrogenase A (LDHA), was not altered by VEGF stimulation in U87/EGFR (Fig 5D) and A172 cells (Fig EV5D). However, VEGF strongly upregulated PFKP expression in several GBM cells, including U87/EGFR (Fig 5D), LN18, U373MG, T98G, and LN229 cells (Fig EV5E) and marginally induced PFKL expression (Fig 5D and EV5D). The increased expression of PFKP plays a role in the regulation of PFK activity (Lee *et al*., 2017). As expected, VEGF stimulation enhanced total PFK enzyme activity in U87/EGFR (Fig 5E) and A172 cells (Fig EV5F).

**Figure 5.**
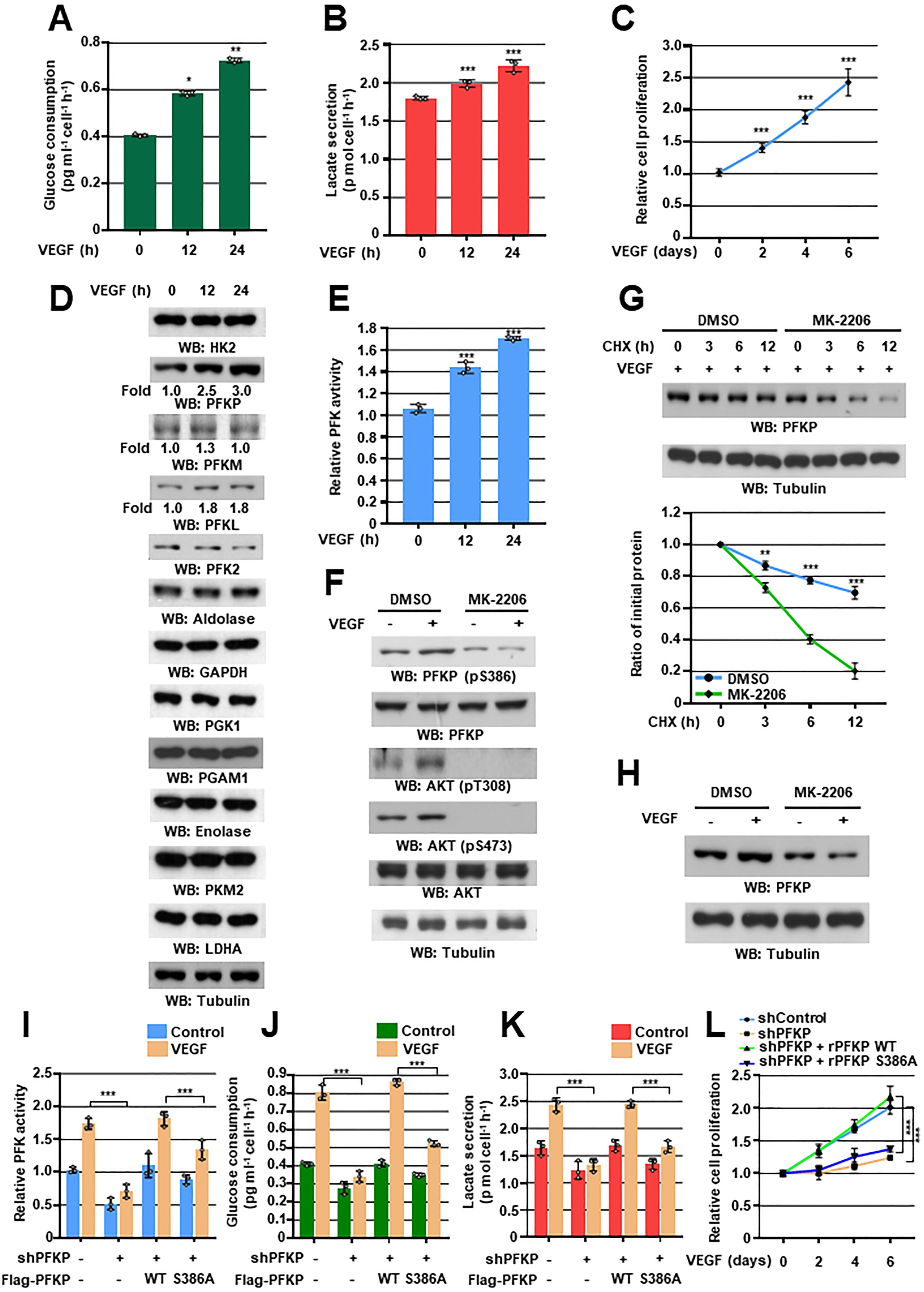
VEGF induces PFKP expression, PFK enzyme activity, aerobic glycolysis, and proliferation in GBM cells. WB and qRT-PCR were performed with the indicated primers and antibodies, respectively (**D, F-H**). **A** and **B** Serum-starved U87/EGFR cells were treated with VEGF (20 ng/mL). Glucose consumption (**A**) and lactate secretion (**B**) were analyzed. **C** U87/EGFR cells in 0.1% serum medium were treated with VEGF (20 ng/mL) and then WST-8 assay was performed. **D** and **E** Serum-starved U87/EGFR cells were treated with VEGF (20 ng/mL). The indicated protein expression levels (**D**) and PFK enzymatic activity (**E**) were measured. **F** Serum-starved U87/EGFR cells were pretreated with DMSO or MK-2206 (5 μM) for 1 h and then stimulated with VEGF (20 ng/mL) for 30 min. **G** Serum-starved U87/EGFR cells were pretreated with VEGF (20 ng/mL) for 1 h and then treated with cycloheximide (CHX;100 μg/mL) in the presence of DMSO or MK-2206 (5 μM). The quantification of PFKP levels relative to tubulin is shown (bottom panel). **H** Serum-starved U87/EGFR cells were pretreated with DMSO or MK-2206 (5 μM) for 1 h and then stimulated with or without VEGF (20 ng/mL) for 24 h. **I-K** U87/EGFR cells with or without the expression of PFKP shRNA and with or without the reconstituted expression of WT Flag-rPFKP or Flag-rPFKP S386A were cultured in serum-free DMEM with or without VEGF (20 ng/mL) for 24 h. PFK enzymatic activity (**I**), glucose consumption **(J),** and lactate secretion **(K)** were analyzed. **L** U87/EGFR cells with or without the expression of PFKP shRNA and with or without the reconstituted expression of WT Flag-rPFKP or Flag-rPFKP S386A were cultured in 0.1% serum medium with or without VEGF (20 ng/mL) and then WST-8 assay was performed. The data represent the mean ± SD of three independent experiments (**A-C, E, G, I-L**). *\*P* < 0.05; *\*\*P* < 0.01; *\*\*\*P* < 0.001, based on the Student’s t-test or one-way ANOVA with Tukey’s post hoc test.

We previously reported that AKT phosphorylated PFKP at S386 to inhibit TRIM21-mediated degradation of PFKP, resulting in increased PFKP expression (Lee *et al*., 2017). VEGF successfully induced AKT phosphorylation and PFKP S386 phosphorylation in U87/EGFR cells (Fig 5F) and A172 cells (Fig EV5G), which was inhibited by pretreatment with the AKT inhibitor MK-2206. Furthermore, the VEGF-induced half-lives of endogenous PFKP (Fig 5G and EV5H) and PFKP expression (Fig 5H and EV5I) were largely decreased by pretreatment with MK-2206 in U87/EGFR cells and A172 cells. To investigate the role of PFKP S386 phosphorylation in VEGF-induced PFK activity, aerobic glycolysis, and proliferation, we depleted endogenous PFKP in U87/EGFR and A172 cells and reconstituted the expression of WT Flag-rPFKP or Flag-rPFKP S386A in these cells. The depletion of PFKP largely impaired VEGF-induced PFK activity (Fig 5I and EV5J), glucose consumption (Fig 5J and EV5K), lactate production (Fig 5K and EV5L), and proliferation (Fig 5L and EV5M), and this inhibition was rescued by the expression of WT rPFKP, but not by the expression of rPFKP S386A mutant in U87/EGFR cells (Fig 5I-5K) and A172 cells (Fig EV5J-EV5L). Taken together, these results indicate that VEGF induces PFKP expression in a PFKP S386 phosphorylation-dependent manner, leading to enhanced PFK enzyme activity, aerobic glycolysis, and proliferation of GBM cells.

## Discussion

Recent studies have shown that some metabolic enzymes and metabolites in cancer cells have been found to have noncanonical or nonmetabolic functions, which are distinct from their original roles in metabolism, and that these play critical roles in a wide spectrum of instrumental cellular activities, including metabolism, and gene expression (Xu *et al*., 2021). Thus, metabolism can be connected in complex ways to multiple cellular processes in human cancer. Here, we demonstrated that EGFR-phosphorylated PFKP Y64 induces VEGF expression directly through AKT activation-mediated β-catenin S552 phosphorylation and indirectly through AKT/SP1-mediated *HIF-1α* transcriptional expression, thereby enhancing blood vessel formation of GBM tumors (Fig 6). Furthermore, VEGF induced the phosphorylation of PFKP S386, resulting in increased PFKP expression, PFK enzyme activity, aerobic glycolysis, and proliferation in GBM cells (Fig 6). Our strong evidence highlights that the nonmetabolic function of PFKP induces angiogenesis and that angiogenesis-independent role of VEGF induces aerobic glycolysis and cancer proliferation, which are mediated by reciprocal regulation between PFKP and VEGF to promote GBM tumor growth (Fig 6).

**Figure 6.**
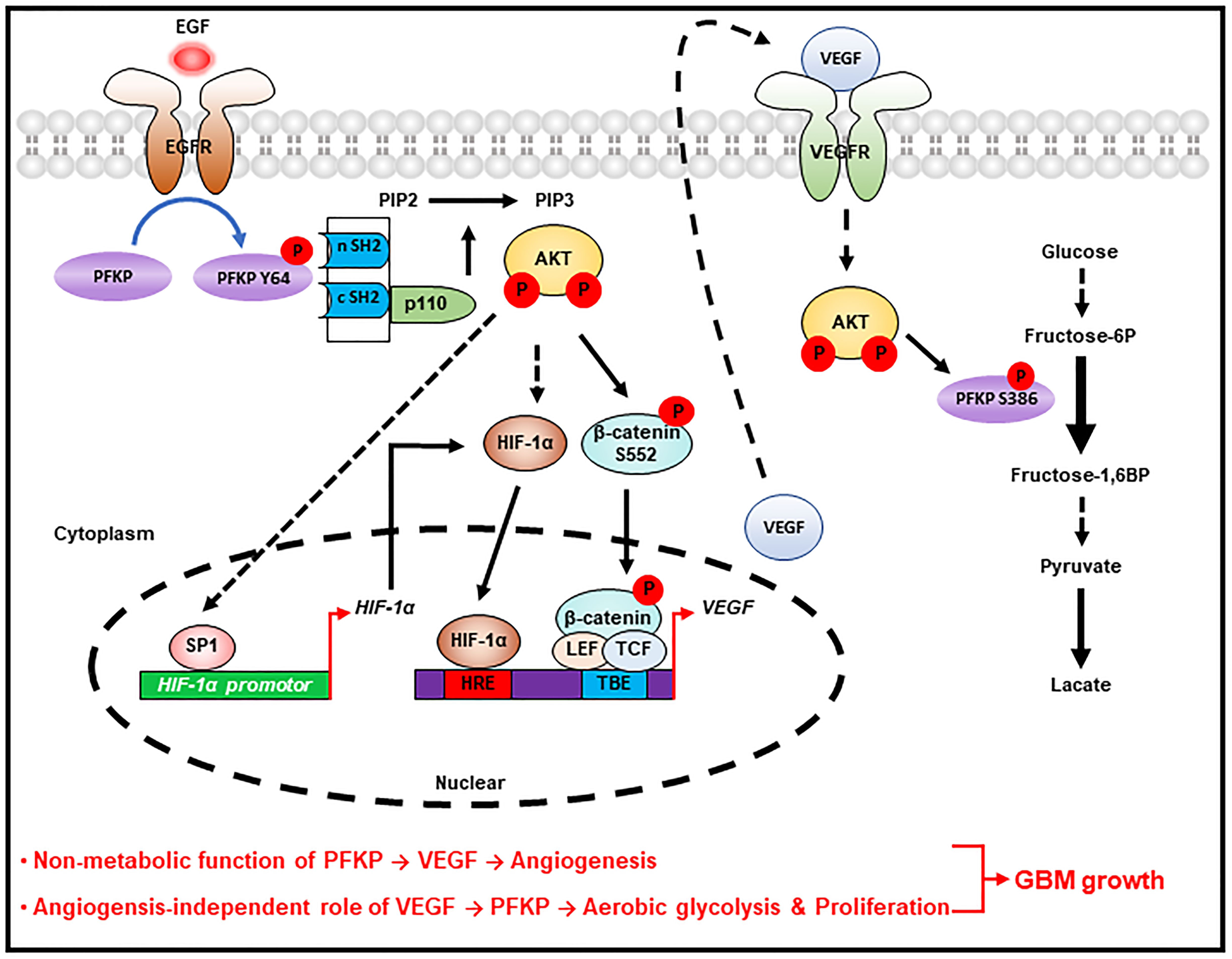
Reciprocal regulation between PFKP and VEGF to promote GBM tumor growth. A schematic of the proposed reciprocal regulation that occurs between PFKP and VEGF in regulating GBM tumor growth. HRE; Hypoxia response element, TBE; TCF binding element.

Studies have shown that HIF-1α accumulation results from intrinsic factors, including gain-of-function of oncoproteins (such as EGFR) and/or loss-of-function of tumor suppressors (such as PTEN) (Laughner *et al*., 2001; Maity *et al*., 2000). We and others (Laughner *et al*., 2001; Maity *et al*., 2000) have shown that increased activity of EGFR, which correlates with poor clinical prognosis in many types of tumor (Avraham & Yarden, 2011), is associated with increased HIF-1α expression in a manner distinct from that mediated by hypoxia (Maxwell *et al*, 1999). Growth factor stimulation induces translation of HIF-1α through activation of the phosphoinositol-3-kinase (PI3K)/AKT/ mammalian target of rapamycin (mTOR) pathway (Guertin & Sabatini, 2007; Page *et al*, 2002). We could not exclude the possibility that PFKP regulates HIF-1α translation, because depletion of PFKP resulted in inhibited AKT-dependent mTOR S2448 phosphorylation (data not shown), that reflects mTOR activity required for HIF-1α translation (Guertin & Sabatini, 2007; Page *et al*., 2002) in response to EGFR activation. Taken together, our data and that of others strongly suggest that the PI3K/AKT pathway could control HIF-1α expression at both the transcriptional and translational levels under normoxic conditions, and our data defines a central role of PFKP Y64 phosphorylation in EGFR activation-induced HIF-1α upregulation in GBM cells.

In earlier work, our *in vivo* tumor xenograft experiments revealed that PFKP was required for brain tumor growth (Lee *et al*., 2018; Lee *et al*., 2017), and in this study, we found that PFKP abrogation results in decreased blood vessel formation *in vivo.* Thus, we evaluated a close association between PFKP and VEGF expression. In this study, we found that blockade of HIF-1α transactivation resulted in marginally inhibited EGFR activation-induced VEGF expression. In addition, exogenous HIF-1α expression could not fully rescue the inhibitory effect of PFKP Y64F mutant expression on the EGFR activation-induced VEGF expression, suggesting that another pathway (i.e. a HIF-1α-independent mechanism) also plays a role in regulating PFKP Y64 phosphorylation-induced VEGF expression in response to EGFR activation. Recently, we have reported that the levels of activated nuclear β-catenin are positively correlated with glioma grades (Du *et al*, 2020) and that PFKP plays an instrumental role in EGFR activation-induced β-catenin transactivation in GBM cells (Lee *et al*., 2020). Consistent with other report that β-catenin directly induces VEGF expression in human colon cancer cells (Easwaran *et al*., 2003), we found that β-catenin activation by AKT-mediated β-catenin S552 phosphorylation plays an important function in EGFR activation-induced *VEGF* transcription in a PFKP Y64 phosphorylation-dependent manner in GBM. Taken together, these observations demonstrate that transcriptional activities of both HIF-1α and β-catenin are required for EGFR activation-induced VEGF expression in GBM cells, in which PFKP Y64 phosphorylation plays a central role. Our *in vivo* data also provides strong evidence that EGFR-phosphorylated PFKP Y64 induces expression levels of HIF-1α, β-catenin S552 phosphorylation, and VEGF, leading to enhanced vascularization of tumors in the GBM xenograft. The clinical significance of these findings was evidenced by the positive correlation between PFKP Y64 phosphorylation and HIF-1α expression, β-catenin S552 phosphorylation, and VEGF expression in human GBM specimens.

The secreted VEGF acts not only on vascular endothelial cells *via* paracrine (Apte *et al*., 2019) for angiogenesis-dependent tumor growth, but also on tumor cells *via* autocrine for angiogenesis-independent tumor growth. Co-expression of VEGF and VEGF receptors (VEGFRs) is commonly observed in a variety of tumor cells, including GBM (Knizetova *et al*, 2008), which enables VEGF/VEGFRs autocrine signaling within a tumor mass to regulate proliferation and tumor growth (Frank *et al*, 2011; Knizetova *et al*., 2008; Lichtenberger *et al*, 2010; Masood *et al*, 2001; Ohba *et al*, 2014; Wu *et al*, 2006). In this study, we provided important evidence that VEGF causes metabolic alterations mainly through PFKP upregulation in GBM cells. We previously reported that AKT-mediated phosphorylation of PFKP at S386 inhibits TRIM21-mediated polyubiquitylation and degradation of PFKP (Lee *et al*., 2017). Here, we found that VEGF signaling activated AKT to induce PFKP S386 phosphorylation, resulting in enhanced PFKP expression, PFK enzyme activity, and aerobic glycolysis, and subsequent proliferation in GBM cells. These collective findings suggest that the VEGF is required for cell-autonomous tumor cell proliferation at least in part in a PFKP expression-dependent manner in human GBM cells.

In summary, our study represents the first demonstration of a reciprocal action between PFKP and VEGF signaling, which may contribute to their overexpressing each other within a tumor mass. This finding highlights novel mechanisms underlying a nonmetabolic function of glycolytic enzyme PFKP-mediated angiogenesis and angiogenesis-independent functions of VEGF-mediated metabolic regulation, which are integrated and mutually regulated to promote GBM tumor growth. These findings underscore that cancer cells’ fundamental biological processes, metabolism, and other cellular activities, are integrated and mutually regulated in promoting tumor development.

## Materials and Methods

### Materials

Rabbit polyclonal antibodies recognizing PFKP (12746, 1:1,000 for immunoblotting), β-catenin (pS552, 9566; 1:1,000 for immunoblotting, 1:300 for immunohistochemistry), EKR1/2 (pT202/pY204, 9101; 1:1,000 for immunoblotting), ERK1/2 (9102; 1:1,000 for immunoblotting), AKT (pS473, 4060; 1:1,000 for immunoblotting), AKT (9272; 1:1,000 for immunoblotting), p38 (pT180/pY182, 9211; 1:1,000 for immunoblotting), p38 (9212; 1:1,000 for immunoblotting), JNK (pT183/pY185, 9251; 1:1,000 for immunoblotting), c-Jun (pS63, 2361; 1:1,000 for immunoblotting), c-Jun (9165; 1:1,000 for immunoblotting), IκBα (pS32/36, 9246; 1:1,000 for immunoblotting), IκBα (9242; 1:1,000 for immunoblotting), p65 (pS536, 3033; 1:1,000 for immunoblotting), p65 (8242; 1:1,000 for immunoblotting), PGK1 (68540, 1:1000 for immunoblotting), and PKM2 (3198, 1:1000 for immunoblotting) were purchased from Cell Signaling Technology (Danvers, MA). Mouse monoclonal antibodies for VEGF (C-1, sc-7269; 1:500 for immunoblotting, 1:100 for immunohistochemistry), SP1 (1C6, sc-420; 1:500 for immunoblotting), PFKL (A-6, sc-393713, 1:500 for immunoblotting), PFKM (sc-67028, 1:1,000 for immunoblotting), HK2 (sc-130858, 1:500 for immunoblotting), LDHA (sc-137243, 1:500 for immunoblotting), GAPDH (sc-47724, 1:500 for immunoblotting), PFK2 (sc-377416, 1:500 for immunoblotting), enolase (sc-271384, 1:500 for immunoblotting), aldolase (sc-390733, 1:500 for immunoblotting), and mithramycin A (SC-200909) were purchased from Santa Cruz Biotechnology (Santa Cruz, CA). Rabbit polyclonal EGFR antibody (pY869, 11229; 1:1,000 for immunoblotting) was purchased, and rabbit polyclonal PFKP antibody (pY64; 5 mg/mL used for immunohistochemical staining) (Lee *et al*., 2018) was customized from Signalway Antibody (College Park, MD). Mouse monoclonal antibody PGAM1 (GTX629754, 1:500 for immunoblotting) was purchased from GeneTex (Irvine, CA). Rabbit polyclonal antibody that recognizes PFKP (pS386; 1:1,000 for immunoblotting) (Lee *et al*., 2017) was customized from Signalway Biotechnology (Pearland, TX). Rabbit monoclonal antibody for HIF-1α (EP1215Y, ab51608; 1:1,000 for immunoblotting, 1:300 for immunohistochemistry) and rabbit polyclonal antibody for CD31 (ab28364; 1:300 for immunohistochemistry) were purchased from Abcam (Cambridge, MA). Mouse monoclonal antibody for EGFR (610016; 1:1,000 for immunoblotting) was purchased from BD Biosciences (San Jose, CA). Mouse monoclonal antibodies for FLAG (clone M2, F3165; 1:5,000 for immunoblotting), HA (H6908; 1:5,000 for immunoblotting), tubulin (clone B-5-1-2, T6074; 1:5,000 for immunoblotting), human recombinant EGF (E9644), hygromycin B (H3274), puromycin (P8833), cycloheximide (66-81-9), and actinomycin D (A1410) were purchased from Sigma (St. Louis, MO). G418 (4727878001) was purchased from Roche (Basel, Switzerland). LY294002 (L-7988), SP600125 (S-7979), PD98059 (P-4313), and SB203580 (S-3400) were purchased from LC Laboratories (Woburn, MA). NF-κB inhibitor (481412) was purchased from EMD Biosciences (San Diego, CA). MK-2206 (S1078) and HIF-1α inhibitor (S7612) was purchased from Selleck Chemicals (Houston, TX). Recombinant human VEGF165 (100-20) was purchased from Peprotech Korea (Seoul, Republic of Korea).

### Cell culture

U373MG GBM cells, A431 human epidermoid carcinoma cells, and MDA-MB-231 human breast carcinoma cells were purchased from the Korean Cell Line Bank (KCLB; Seoul, Republic of Korea). LN18, T98G, A172, and LN229 GBM cells were kindly provided by Dr. Hyunggee Kim (Korea University, Seoul, Republic of Korea). U87 and U251 GBM cells were kindly provided by Dr. Kyu Heo (Dongnam Institute of Radiological and Medical Sciences, Busan, Republic of Korea). All cells were routinely tested for mycoplasma. These tumor cells were maintained in Dulbecco’s modified Eagle’s medium (DMEM) supplemented with 10% fetal bovine serum (Capricon, USA).

### Transfection

Cells were plated at a density of 4 × 10^5^ per 60-mm dish or 1 × 10^5^ per well of a 6-well plate, 18 h before transfection. Transfection was performed using Lipofectamine^2000^ transfection reagent (ThermoFisher; Pittsburgh, PA) according to the manufacturer’s instructions. SP1 siRNA (116546) was purchased from ThermoFisher (Pittsburgh, PA). Transfection of SP1 siRNA was performed using Lipofectamine^TM^ RNAiMAX transfection reagent (ThermoFisher; Pittsburgh, PA) according to the manufacturer’s instructions.

### Quantitative real-time PCR analysis

Total RNA was prepared from tumor cells using an RNeasy Mini kit (Qiagen; Valencia, CA) according to the manufacturer’s instructions, and cDNA was synthesized from 2 μg of total RNA by reverse transcriptase (Superscript II Preamplification System, Gibco-BRL; Gaithersburg, MD). Real-time PCR was performed on an ABI Prism 7500 sequence detection system using a SYBR® Green PCR Master Mix (Applied Biosystems; Foster City, CA) and following the manufacturer’s protocols. The ABI 7500 sequence detector was programmed with the following PCR conditions: 40 cycles of 15-s denaturation at 95°C and 1-minute amplification at 60°C. All reactions were run in triplicate and normalized to the housekeeping gene *HPRT*. The evaluation of relative differences of PCR results was calculated using the comparative cycle threshold (CT) method. The following primer pairs were used for quantitative real-time PCR: human *HIF-1α*, 5′-CATAAAGTCTGCAACATGGAAGGT-3′ (forward) and 5′-ATTTAGTGGGTGAGGAATGGGTT-3′ (reverse); human *VEGF*, 5′-TGCAGATTATGCGGATCAAACC-3′ (forward) and 5′-TGCATTCACATTTGTTGTGCTGTAG-3′ (reverse); and human *HPRT*, 5′-CATTATGCTGAGGATTTGGAAAGG-3′ (forward) and 5′-CTTGAGCACACAGAGGGCTACA-3′ (reverse).

### Western blot analysis

Extraction of proteins from cultured cells was performed using a lysis buffer (50 mM Tris-HCl, [pH 7.5], 0.1% SDS, 1% Triton X-100, 150 mM NaCl, 1 mM DTT, 0.5 mM EDTA, 100µM sodium orthovanadate, 100µM sodium pyrophosphate, 1 mM sodium fluoride, and proteinase inhibitor cocktail). Cell extracts were clarified via centrifugation at 13,400 g, and the supernatants (1.5 mg protein/mL) were subjected to immunoblot analysis with corresponding antibodies. Band intensity was quantified using ImageJ 1.53e software (National Institutes of Health). Each experiment was repeated at least three times.

### Luciferase reporter assay

The tumor cells were co-transfected with pGL3 empty vector, pGL3-HRE-luciferase plasmid containing five copies of HREs from human VEGF gene, pGreenFire1-mCMV, or pGreenFire1-SP1 (System Biosciences; Palo Alto, CA) and pRL-TK vector (as an inner control that contains Renilla luciferase sequences (Promega; Madison, WI)) using Lipofectamine^2000^ transfection reagent (ThermoFisher; Pittsburgh, PA) according to the manufacturer’s instructions, and then grown under different experimental conditions. After incubation, firefly and Renilla luciferase activities were measured using a Dual-Luciferase® Reporter Assay System (Promega; Madison, WI), and the ratio of firefly/Renilla luciferase was determined.

### Chromatin Immunoprecipitation (ChIP) assay

A ChIP assay was performed using the SimpleChIP Enzymatic Chromatin IP kit (9003; Cell Signaling Technology). Chromatin prepared from 2 × 10^6^ cells (in a 10-cm dish) was used to determine the total DNA input and was incubated overnight with SP1 antibody or with normal mouse IgG. The immunoprecipitated chromatin was detected using real-time PCR. The PCR primer sequences for the *HIF-1α* promoter were SP1 #P1, 5′-CGAGGCGAAGTCTGCTTTTT-3′ (forward) and 5′-TCCTACTCTTGGTGCAGTAATG-3′ (reverse); SP1 #P2, 5′-TCGCTCGCCATTGGATCTCG-3′ (forward) and 5′-GCGCGCGGGGAGGGGAGAGG-3′ (reverse); and SP1 #P3, 5′-CCCCCTCTCCCCTCCCCGCG-3′ (forward) and 5′-GAGGAGCTGAGGCAGCGTCA-3′ (reverse).

### Measurement of glucose consumption and lactate production

Cells were seeded in culture dishes, and the medium was changed after 12 hours with non-serum containing DMEM. The cells were incubated for indicated periods of time, and the culture medium was collected for the measurement of glucose and lactate concentrations. Glucose levels were determined using a glucose (GO) assay kit (Sigma-Aldrich). Glucose consumption was calculated as the difference in glucose concentration between the collected culture medium and DMEM. Absorbance was recorded at 540 nm at room temperature in a 96-well plate. Lactate levels were determined using a lactate assay kit (Eton Bioscience, San Diego, CA). Absorbance was recorded at 570 nm at room temperature in a 96-well plate. All results were normalized to the final cell number.

### Measurement of PFK activity

PFK activity was assessed using a phosphofructokinase (PFK) activity colorimetric assay kit (BioVision, Milpitas, CA). The reaction was performed using cell lysate (3 µg) in 100 µL of reaction buffer, which was prepared according to the kit instructions. Absorbance was recorded at 450 nm at 37°C in a 96-well plate.

### Cell proliferation assay

Cells were plated at a density of 1 × 10^3^ per 96-well plate. Cells were treated VEGF (20 ng/mL) for the indicated periods days in DMEM with 0.1% serum. Cell proliferation was measured by Quanti-Max^TM^ WST-8 cell viability assay kit (BIOMAX, Republic of Korea). The reaction was performed by treating Quanti-Max^TM^ (10 µL per well) in culture media, and then incubated for 0.5 ∼ 2 h at 37°C. Absorbance was recorded at 450 nm at 37°C.

### Intracranial implantation of GBM cells in mice and histologic evaluation

We injected U87/EGFRvIII GBM cells with or without modulation of PFKP expression or an active AKT1 mutant, intracranially into 4-week-old male athymic nude mice (five mice/group), as described previously (Lee *et al*., 2018). The mice were euthanized 2 weeks after the GBM cells were injected. The brain of each mouse was harvested, fixed in 4% formaldehyde, and embedded in paraffin. After that, histological sections (5 μm) were prepared. The sections were stained with Mayer’s hematoxylin and subsequently with eosin (H&E) (Biogenex Laboratories, San Ramon, CA). Afterward, the slides were mounted with Universal Mount (Research Genetics Huntsville, AL). Tumor formation and phenotype were determined by histological analysis of H&E-stained sections. Tumor volume was calculated by 0.5 × L × W2 (L, length; W, width). All of the mice were housed in the in the Animal Central Laboratory of West China the Second Hospital (Chengdu, Sichuan) animal facility, and all experiments were performed in accordance with relevant institutional and national guidelines and regulations approved by the Institutional Animal Care and Use Committee at State Key Laboratory of Oral Diseases, Sichuan University.

### IHC analysis and scoring

An IHC analysis was conducted using paraffin-embedded tissue sections. The expression of HIF-1α, β-catenin S552 phosphorylation, VEGF, and CD31 was detected with a VECTASTAIN Elite ABC kit (Vector Laboratories); tissue sections were then incubated with 3,3′-diaminobenzidine (Vector Laboratories), and the nuclei were stained with hematoxylin. Six randomly chosen fields per slide were analyzed and averaged.

The human GBM samples and clinical information were from the Chinese Glioma Genome Atlas (CGGA, http://www.cgga.org.cn). This study was approved by the Ethics Committee of Capital Medical University (China), and written informed consents were obtained from all patients. The tissue sections from 25 paraffin-embedded human GBM specimens were stained with antibodies against PFKP Y64 phosphorylation, HIF-1α, β-catenin S552 phosphorylation, VEGF, or non-specific immunoglobulin as a negative control. We quantitatively scored the tissue sections according to the percentage of positive cells and staining intensity, as previously defined (Lee *et al*., 2018). We assigned the following proportion scores: 0 if 0% of the tumor cells showed positive staining, 1 if 0.1% to 1% of cells were stained, 2 if 1.1% to 10% stained, 3 if 11% to 30% stained, 4 if 31% to 70% stained, and 5 if 71% to 100% stained. We rated the intensity of staining on a scale of 0 to 3: 0, negative; 1, weak; 2, moderate; and 3, strong. We then combined the proportion and intensity scores to obtain a total score (range, 0–8), as described previously (Lee *et al*., 2018). The use of human glioblastoma samples and the clinical parameters was approved by the Institutional Review Board at Capital Medical University in Beijing, China.

### TCGA analyses

TCGA data were downloaded from cBioPortal (https://www.cbioportal.org). Correlation analysis between two genes was done with *Pearson*’s correlation analysis. *P*-value indicates significance of correlation.

### Statistical analysis

All quantitative data are presented as the mean ± SD of at least three independent experiments. A 2-group comparison was conducted using the 2-sided, 2-sample Student’s t-test. A simultaneous comparison of >2 groups was conducted using one-way ANOVA followed by Tukey’s post hoc tests. The SPSS statistical package (version 12; SPSS Inc.) was used for the analyses. Values of P < 0.05 were considered to indicate statistically significant differences.

## Acknowledgments

The pGL3-HRE-luciferase plasmid was kindly provided by Dr. You Mie Lee (Kyungpook National University, Republic of Korea). This work was supported by the National Research Foundation of Korea (NRF) grant funded by the Korean government (MIST; 2020R1C1C1011350).

## Author’s contributions

JSL, YS, LD, and J-HL designed the study. JSL, YS, SMJ, SHP, CZ, Y-YP, RL, JL, W-SC, and J-HL performed the experimental work. JSL, YS, Y-YP, RL, JL, W-SC, LD, and J-HL performed data analyses. JSL, YS, Y-YP, JL, W-SC, LD, and J-HL produced the main draft of the text and the figures. All authors critically revised and approved the final manuscript.

## Data availability

All data generated or analyzed during this study are included in this published article.

## Expanded View Figure legends

**Figure EV1 (related to Figure 1).**
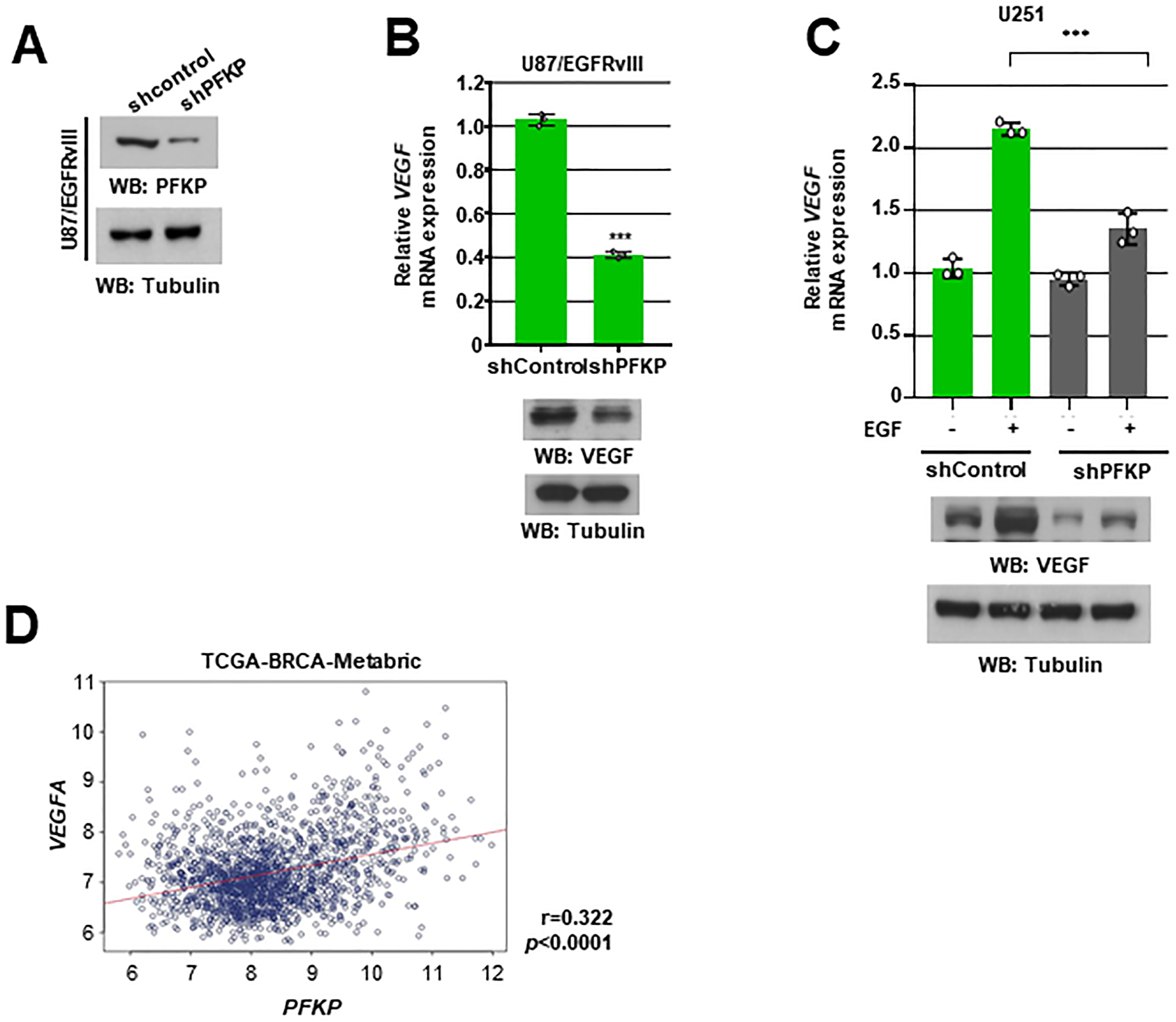
PFKP depletion in GBM cells results in impaired EGFR activation-induced VEGF expression. WB and qRT-PCR were performed with the indicated primers and antibodies, respectively (**A-C**). **A** U87/EGFRvIII cells were transfected with shRNA against PFKP. **B** Expression levels of VEGF in the U87/EGFRvIII cells stably expressing control shRNA or PFKP shRNA. **C** Serum-starved U251 cells with or without PFKP depletion were treated with EGF (100 ng/mL) for 12 h. **D** TCGA analysis of *PFKP* and *VEGF* mRNA expression in breast tumor specimens (n = 1094). The data represent the mean ± SD of three independent experiments (**B, C**). \*\*\**P* < 0.001, based on the Student’s t-test.

**Figure EV2 (related to Figure 1).**
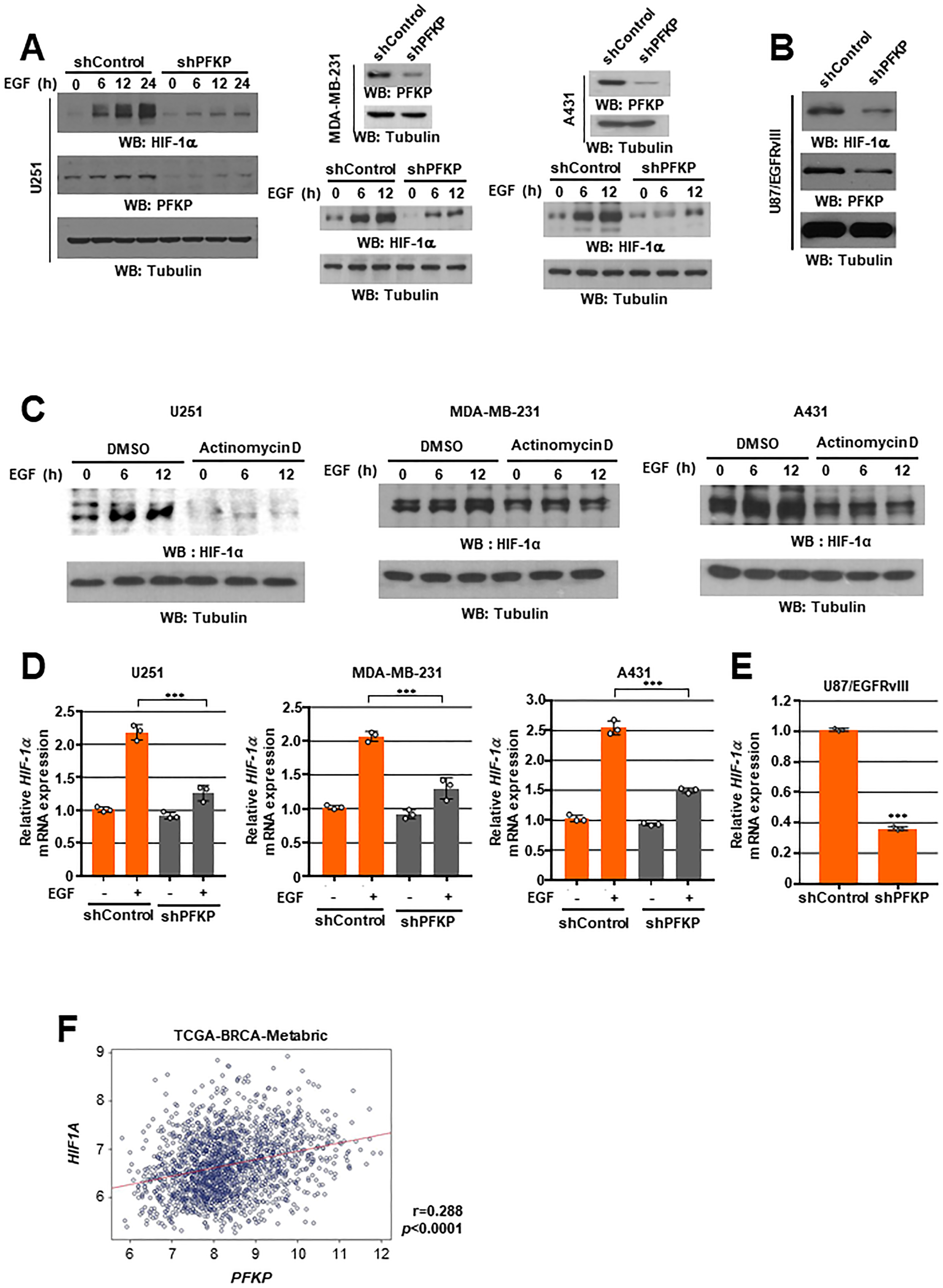
PFKP expression is required for EGFR activation-induced HIF-1α expression and transactivation. WB and qRT-PCR were performed with the indicated primers and antibodies, respectively (**A-E**). **A** Serum-starved U251, MDA-MB-231, and A431 cells with or without PFKP depletion were treated with EGF (100 ng/mL). **B** U87/EGFRvIII were stably expressed with control shRNA or PFKP shRNA. **C** Serum-starved U251, MDA-MB-231, and A431 cells were pretreated DMSO or actinomycin D (1 μg/mL) for 1 h and then stimulated with or without EGF (100 ng/mL). **D** Serum-starved U251, MDA-MB-231, and A431 cells stably expressed control shRNA or shPFKP were treated with or without EGF (100 ng/mL) for 12 h. **E** U87/EGFRvIII cells were stably expressed with control shRNA or PFKP shRNA. **F** TCGA analysis of *PFKP* and *HIF-1α* mRNA expression in breast tumor specimens (n = 1094). The data represent the mean ± SD of three independent experiments (**D, E**). \*\*\**P* < 0.001, based on the Student’s t-test.

**Figure EV3 (related to Figure 2).**
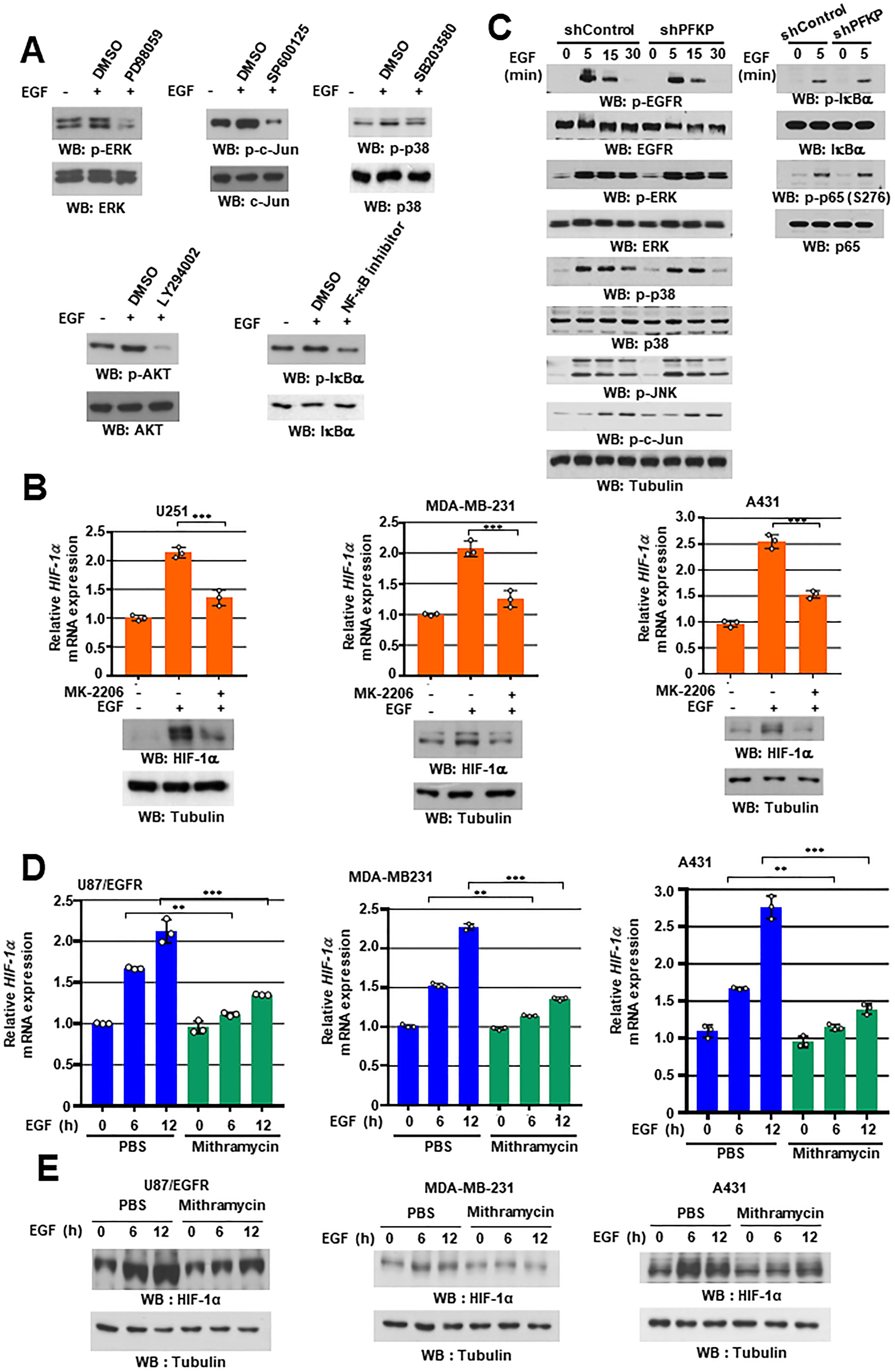
PFKP Y64 phosphorylation induces EGFR activation-enhanced *HIF-1α* transcriptional expression through SP1 transactivation. WB and qRT-PCR were performed with the indicated primers and antibodies, respectively (**A-E**). **A** Serum-starved U87/EGFR cells were pretreated with the indicated inhibitors for 1 h and then stimulated with or without EGF (100 ng/mL) for 30 min. **B** Serum-starved U251, MDA-MB-231, and A431 cells were pretreated with DMSO or MK-2206 (5 μM) for 1h and then stimulated with or without EGF (100 ng/mL) for 12 h. **C** Serum-starved U87/EGFR cells stably expressing control shRNA or shPFKP were treated with or without EGF (100 ng/mL). **D** and **E** U87/EGFR, MDA-MB-231, A431 cells were pretreated with PBS or mithramycin (500 nM) for 1 h and then stimulated with or without EGF (100 ng/mL). The data represent the mean ± SD of three independent experiments (**B, D**). \*\**P* < 0.01; \*\*\**P* < 0.001, based on the Student’s t-test.

**Figure EV4 (related to Figure 3).**
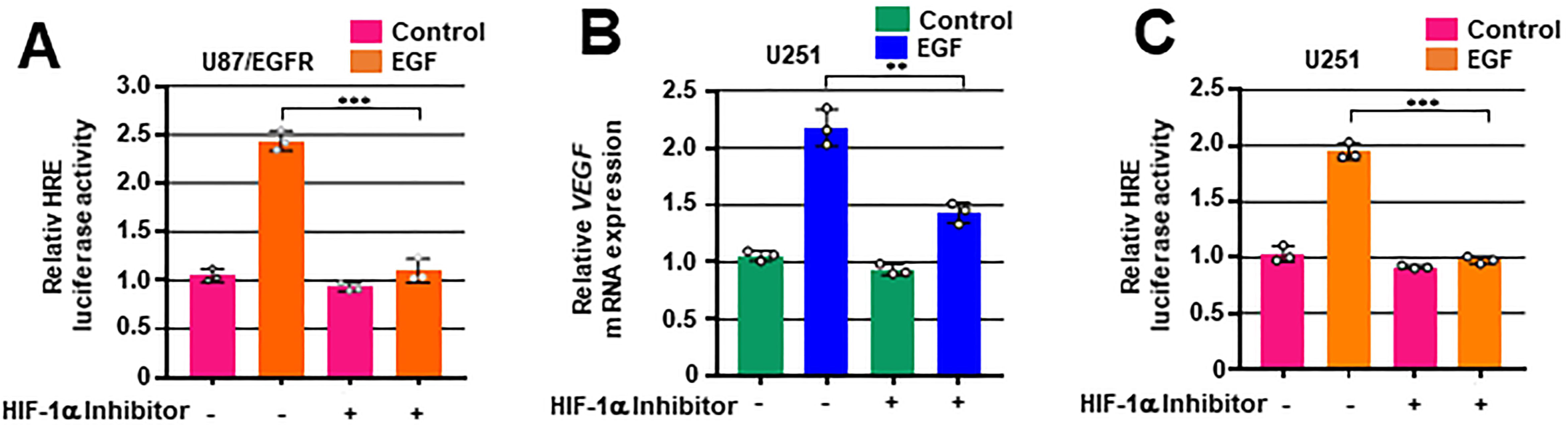
PFKP Y64 phosphorylation induces VEGF expression through HIF-1 α expression and β-catenin Ser552 phosphorylation in response to EGFR activation. **A** and **C** Serum-starved U87/EGFR **(A)** and U251 **(C)** cells were co-transfected with luciferase reporter plasmids (pGL3-HRE-luciferase) and the Renilla control plasmid. The cells were pretreated with DMSO or HIF-1α inhibitor (10 μM) for 1 h and then stimulated with or without EGF (100 ng/mL) for 12 h. Luciferase activity was measured. **B** Serum-starved U251 cells were pre-treated DMSO or HIF-1α inhibitor (10 μM) for 1h and then stimulated with or without EGF (100 ng/mL) for 12 h. The mRNA expression level of VEGF was determined by real-time PCR. The data represent the mean ± SD of three independent experiments. \*\**P* < 0.01; \*\*\**P* < 0.001, based on the Student’s t-test.

**Figure EV5 (related to Figure 5).**
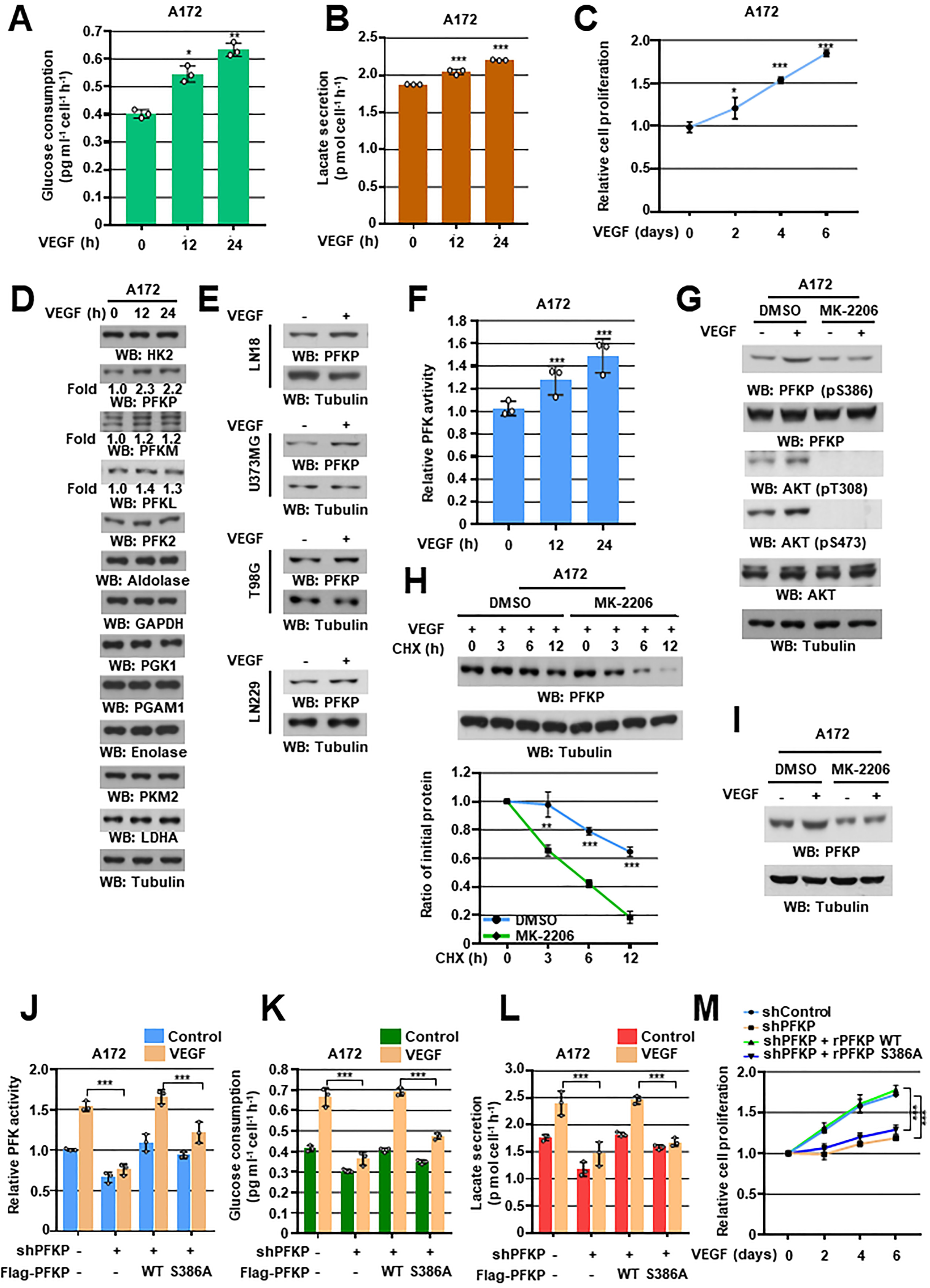
VEGF induces PFKP expression, PFKP expression, PFK enzyme activity, aerobic glycolysis, and proliferation in GBM cells. WB and qRT-PCR were performed with the indicated primers and antibodies, respectively (**D, E, G-I**). **A** and **B** Serum-starved A172 cells were treated with VEGF (20 ng/mL). Glucose consumption (**A**) and lactate secretion (**B**) were analyzed. **C** A172 cells in 0.1% serum medium were treated with VEGF (20 ng/mL) and then WST-8 assay was performed. **D** and **F** Serum-starved A172 cells were treated with VEGF (20 ng/mL). The indicated protein expression levels (**D**) and PFK enzymatic activity (**F**) were measured. **E** Serum-starved LN18, U373MG, T98G, and LN229 cells were treated with or without VEGF (20 ng/ml) for 12h. **G** Serum-starved A172 cells were pretreated with DMSO or MK-2206 (5 μM) for 1 h and then stimulated with VEGF (20 ng/mL) for 30 min. **H** Serum-starved A172 cells were pretreated with VEGF (20 ng/mL) for 1 h and then treated with cycloheximide (CHX;100 μg/mL) in the presence of DMSO or MK-2206 (5 μM). The quantification of PFKP levels relative to tubulin is shown (bottom panel). **I** Serum-starved A172 cells were pretreated with DMSO or MK-2206 (5 μM) for 1 h and then stimulated with or without VEGF (20 ng/mL) for 24 h. **J-L** A172 cells with or without the expression of PFKP shRNA and with or without the reconstituted expression of WT Flag-rPFKP or Flag-rPFKP S386A were cultured in serum-free DMEM with or without VEGF (20 ng/mL) for 24 h. PFK enzymatic activity (**J**), glucose consumption **(K),** and lactate secretion **(L)** were analyzed. **M** A172 cells with or without the expression of PFKP shRNA and with or without the reconstituted expression of WT Flag-rPFKP or Flag-rPFKP S386A were cultured in 0.1% serum medium with or without VEGF (20 ng/mL) and then WST-8 assay was performed. The data represent the mean ± SD of three independent experiments (**A-C, F, H, J-M**). *\*P* < 0.05; *\*\*P* < 0.01; *\*\*\*P* < 0.001, based on the Student’s t-test or one-way ANOVA with Tukey’s post hoc test.

## References

Apte RS, Chen DS, Ferrara N (2019) VEGF in Signaling and Disease: Beyond Discovery and Development. Cell 176: 1248–1264

Avraham R, Yarden Y (2011) Feedback regulation of EGFR signalling: decision making by early and delayed loops. Nat Rev Mol Cell Biol 12: 104–117

Cancer Genome Atlas Research N (2008) Comprehensive genomic characterization defines human glioblastoma genes and core pathways. Nature 455: 1061–1068

Chuang CW, Pan MR, Hou MF, Hung WC (2013) Cyclooxygenase-2 up-regulates CCR7 expression via AKT-mediated phosphorylation and activation of Sp1 in breast cancer cells. J Cell Physiol 228: 341–348

Du L, Lee JH, Jiang H, Wang C, Wang S, Zheng Z, Shao F, Xu D, Xia Y, Li J et al (2020) beta-Catenin induces transcriptional expression of PD-L1 to promote glioblastoma immune evasion. J Exp Med 217

Easwaran V, Lee SH, Inge L, Guo L, Goldbeck C, Garrett E, Wiesmann M, Garcia PD, Fuller JH, Chan V et al (2003) beta-Catenin regulates vascular endothelial growth factor expression in colon cancer. Cancer Res 63: 3145–3153

Fang D, Hawke D, Zheng Y, Xia Y, Meisenhelder J, Nika H, Mills GB, Kobayashi R, Hunter T, Lu Z (2007) Phosphorylation of beta-catenin by AKT promotes beta-catenin transcriptional activity. J Biol Chem 282: 11221–11229

Feldkamp MM, Lau N, Rak J, Kerbel RS, Guha A (1999) Normoxic and hypoxic regulation of vascular endothelial growth factor (VEGF) by astrocytoma cells is mediated by Ras. Int J Cancer 81: 118–124

Forsythe JA, Jiang BH, Iyer NV, Agani F, Leung SW, Koos RD, Semenza GL (1996) Activation of vascular endothelial growth factor gene transcription by hypoxia-inducible factor 1. Mol Cell Biol 16: 4604–4613

Frank NY, Schatton T, Kim S, Zhan Q, Wilson BJ, Ma J, Saab KR, Osherov V, Widlund HR, Gasser M et al (2011) VEGFR-1 expressed by malignant melanoma-initiating cells is required for tumor growth. Cancer Res 71: 1474–1485

Guertin DA, Sabatini DM (2007) Defining the role of mTOR in cancer. Cancer Cell 12: 9–22

Hanahan D, Weinberg RA (2011) Hallmarks of cancer: the next generation. Cell 144: 646–674

Hung MS, Chen IC, Lin PY, Lung JH, Li YC, Lin YC, Yang CT, Tsai YH (2016) Epidermal growth factor receptor mutation enhances expression of vascular endothelial growth factor in lung cancer. Oncol Lett 12: 4598–4604

Iyer NV, Leung SW, Semenza GL (1998) The human hypoxia-inducible factor 1alpha gene: HIF1A structure and evolutionary conservation. Genomics 52: 159–165

Jain RK, di Tomaso E, Duda DG, Loeffler JS, Sorensen AG, Batchelor TT (2007) Angiogenesis in brain tumours. Nat Rev Neurosci 8: 610–622

Knizetova P, Ehrmann J, Hlobilkova A, Vancova I, Kalita O, Kolar Z, Bartek J (2008) Autocrine regulation of glioblastoma cell cycle progression, viability and radioresistance through the VEGF-VEGFR2 (KDR) interplay. Cell Cycle 7: 2553–2561

Laughner E, Taghavi P, Chiles K, Mahon PC, Semenza GL (2001) HER2 (neu) signaling increases the rate of hypoxia-inducible factor 1alpha (HIF-1alpha) synthesis: novel mechanism for HIF-1-mediated vascular endothelial growth factor expression. Mol Cell Biol 21: 3995–4004

Lee JH, Liu R, Li J, Wang Y, Tan L, Li XJ, Qian X, Zhang C, Xia Y, Xu D et al (2018) EGFR-Phosphorylated Platelet Isoform of Phosphofructokinase 1 Promotes PI3K Activation. Mol Cell 70: 197–210 e197

Lee JH, Liu R, Li J, Zhang C, Wang Y, Cai Q, Qian X, Xia Y, Zheng Y, Piao Y et al (2017) Stabilization of phosphofructokinase 1 platelet isoform by AKT promotes tumorigenesis. Nat Commun 8: 949

Lee JH, Shao F, Ling J, Lu S, Liu R, Du L, Chung JW, Koh SS, Leem SH, Shao J et al (2020) Phosphofructokinase 1 Platelet Isoform Promotes beta-Catenin Transactivation for Tumor Development. Front Oncol 10: 211

Lichtenberger BM, Tan PK, Niederleithner H, Ferrara N, Petzelbauer P, Sibilia M (2010) Autocrine VEGF signaling synergizes with EGFR in tumor cells to promote epithelial cancer development. Cell 140: 268–279

Maity A, Pore N, Lee J, Solomon D, O’Rourke DM (2000) Epidermal growth factor receptor transcriptionally up-regulates vascular endothelial growth factor expression in human glioblastoma cells via a pathway involving phosphatidylinositol 3’-kinase and distinct from that induced by hypoxia. Cancer Res 60: 5879–5886

Masood R, Cai J, Zheng T, Smith DL, Hinton DR, Gill PS (2001) Vascular endothelial growth factor (VEGF) is an autocrine growth factor for VEGF receptor-positive human tumors. Blood 98: 1904–1913

Maxwell PH, Wiesener MS, Chang GW, Clifford SC, Vaux EC, Cockman ME, Wykoff CC, Pugh CW, Maher ER, Ratcliffe PJ (1999) The tumour suppressor protein VHL targets hypoxia-inducible factors for oxygen-dependent proteolysis. Nature 399: 271–275

Mor I, Cheung EC, Vousden KH (2011) Control of glycolysis through regulation of PFK1: old friends and recent additions. Cold Spring Harb Symp Quant Biol 76: 211–216

Moreno-Sanchez R, Rodriguez-Enriquez S, Marin-Hernandez A, Saavedra E (2007) Energy metabolism in tumor cells. FEBS J 274: 1393–1418

Ohba T, Cates JM, Cole HA, Slosky DA, Haro H, Ando T, Schwartz HS, Schoenecker JG (2014) Autocrine VEGF/VEGFR1 signaling in a subpopulation of cells associates with aggressive osteosarcoma. Mol Cancer Res 12: 1100–1111

Page EL, Robitaille GA, Pouyssegur J, Richard DE (2002) Induction of hypoxia-inducible factor-1alpha by transcriptional and translational mechanisms. J Biol Chem 277: 48403–48409

Phillips RJ, Mestas J, Gharaee-Kermani M, Burdick MD, Sica A, Belperio JA, Keane MP, Strieter RM (2005) Epidermal growth factor and hypoxia-induced expression of CXC chemokine receptor 4 on non-small cell lung cancer cells is regulated by the phosphatidylinositol 3-kinase/PTEN/AKT/mammalian target of rapamycin signaling pathway and activation of hypoxia inducible factor-1alpha. J Biol Chem 280: 22473–22481

Pore N, Liu S, Shu HK, Li B, Haas-Kogan D, Stokoe D, Milanini-Mongiat J, Pages G, O’Rourke DM, Bernhard E et al (2004) Sp1 is involved in Akt-mediated induction of VEGF expression through an HIF-1-independent mechanism. Mol Biol Cell 15: 4841–4853

Pouyssegur J, Dayan F, Mazure NM (2006) Hypoxia signalling in cancer and approaches to enforce tumour regression. Nature 441: 437–443

Ravindranath N, Wion D, Brachet P, Djakiew D (2001) Epidermal growth factor modulates the expression of vascular endothelial growth factor in the human prostate. J Androl 22: 432–443

Reisinger K, Kaufmann R, Gille J (2003) Increased Sp1 phosphorylation as a mechanism of hepatocyte growth factor (HGF/SF)-induced vascular endothelial growth factor (VEGF/VPF) transcription. J Cell Sci 116: 225–238

Semenza GL (1998) Hypoxia-inducible factor 1: master regulator of O2 homeostasis. Curr Opin Genet Dev 8: 588–594

Shweiki D, Itin A, Soffer D, Keshet E (1992) Vascular endothelial growth factor induced by hypoxia may mediate hypoxia-initiated angiogenesis. Nature 359: 843–845

Sroka IC, Nagle RB, Bowden GT (2007) Membrane-type 1 matrix metalloproteinase is regulated by sp1 through the differential activation of AKT, JNK, and ERK pathways in human prostate tumor cells. Neoplasia 9: 406–417

Tan NY, Khachigian LM (2009) Sp1 phosphorylation and its regulation of gene transcription. Mol Cell Biol 29: 2483–2488

Wu Y, Hooper AT, Zhong Z, Witte L, Bohlen P, Rafii S, Hicklin DJ (2006) The vascular endothelial growth factor receptor (VEGFR-1) supports growth and survival of human breast carcinoma. Int J Cancer 119: 1519–1529

Xu D, Shao F, Bian X, Meng Y, Liang T, Lu Z (2021) The Evolving Landscape of Noncanonical Functions of Metabolic Enzymes in Cancer and Other Pathologies. Cell Metab 33: 33–50

